# The evolution of sex allocation in metapopulations

**DOI:** 10.1101/2020.05.08.080929

**Authors:** Camille Roux, Charles Mullon, John R. Pannell

**Affiliations:** Univ. Lille, CNRS, UMR 8198 - Evo-Eco-Paleo, F-59000 Lille, France; Department of Ecology and Evolution, Biophore Building, University of Lausanne, 1015 Lausanne, Switzerland

## Abstract

Selection in inbred populations is expected to favor female-biased sex ratios in dioecious populations as a result of local mate competition, a prediction that finds strong support in situations where females have control of the sex ratio. Local mate competition due to inbreeding should also promote female-biased sex allocation in hermaphrodites, with reduced emphasis on the production and dispersal of sperm or pollen relative to that of eggs, ovules or seeds. While inbreeding can be the direct result of the mating system in local populations, it can also be brought about by population turnover in metapopulations with frequent local extinction and recolonization. This effect of population turnover has previously been considered for species with separate sexes. Here, we use both formal analysis and individual-based simulations to explore the effect of population turnover on sex allocation in partially self-fertilizing hermaphroditic metapopulations. Using simulations, we also assess the extent to which different genetic measures of inbreeding and population differentiation predict the equilibrium sex allocation. We find that population turnover can select for strongly female-biased sex allocation in hermaphroditic metapopulations, particularly if among-deme dispersal is low, even where local demes are fully outcrossing. In such situations, *F*_ST_ is a good predictor of the equilibrium sex allocation, and much better than the alternative differentiation measures *G*_ST_ and Jost’s *D*. Our study extends predictions for sex allocation in subdivided populations to hermaphroditic species, and draws attention in general to the power of Wright’s hierarchical inbreeding statistics to predict the sex allocation in metapopulations at equilibrium.

## 1 Introduction

An individual’s sex allocation is the proportion of reproductive resources it invests in each of its sexual functions (ranging between 0 and 1 from male to female extremes). In dioecious species, the sex allocation is the proportion of reproductive resources allocated to sons (i.e., effectively the sex ratio when sons and daughters are equally costly). Düsing (1884) first provided an evolutionary explanation for equal sex ratios, although the explanation is often credited to Fisher (1958). Simply put, because autosomal genes have an equal chance of passing to the next generation through male and female gametes, selection should favor an equal investment in each of these two paths. The corollary is that, in populations with an unequal sex ratio, selection will favor alleles in the minority sex because they must, on average, leave more progeny than those of the majority sex. In such populations, natural selection should thus rapidly bring about equilibration of the sex ratio (Düsing, 1884; Fisher, 1958).

Although equal sex ratios are common, many species produce female-biased sex ratios. Hamilton (1967) explained these ‘extraordinary sex ratios’ largely in terms of the effects of ‘local mate competition’. If mating is not random, and, in particular, if sons compete among themselves for mating with a restricted number of females, selection should favor the production of fewer sons and more daughters (Taylor and Bulmer, 1980). Hamilton (1967)’s prediction is supported by a wide range of empirical observations and represents perhaps the most well-supported prediction of life-history theory (reviewed on page 84 in West, 2009). The prediction is particularly well illustrated by examples where the sons and daughters of a small number of fertilized females hatch together and mate before dispersing, producing inbred progeny (Read et al., 1992). Indeed, the inbreeding coefficient, a measure of the degree to which related individuals mate with one another relative to the case of random mating, is a reliable predictor of the progeny sex ratio (Molbo and Parker, 1996; West et al., 2005; Shuker et al., 2005; Grillenberger et al., 2008; Molbo et al., 2003; Fellowes et al., 1999; Song et al., 2016; Reece et al., 2008).

The effect of local mate competition on the evolution of sex allocation can also be viewed from the perspective of sex-biased dispersal. In the cases just cited, females disperse after mating, carrying the genes of the males they mated with. Biased investment in females is thus selected also because female dispersal effectively contributes to fitness through both sexes. In contrast, if sons disperse more widely than daughters, or, in plants, if pollen is dispersed more widely than seeds (as will be typical for many species; Hamrick and Godt, 1996; Ennos, 1994), selection should favor male-biased sex allocation (de Jong et al., 2002). In general terms, selection should favor allocation bias to the more highly dispersing sex (Guillon and Bottein, 2011; de Jong et al., 2002), although this prediction depends on the details of dispersal, e.g., whether dispersal is within or among demes (Ravigne et al., 2006).

While the above ideas were developed and have been tested principally for dioecious species, they should also apply to hermaphrodites (West, 2009; Charnov, 1982; Brunet, 1992; Charlesworth and Charlesworth, 1981; Schärer, 2009; Yin and Haag, 2019; Cutter, 2015). The sex allocation of hermaphrodites can be viewed in terms of the relative investment made to the production and dispersal of sperms versus eggs or, for seed plants, to that of pollen versus seeds. As is the case for species with separate sexes, selection on the sex allocation of hermaphrodites should favor equal investment in both sexual functions if mating is random and both pollen and seeds are widely dispersed (Lloyd, 1982). Limited dispersal can cause sib competition, and selection should then favor allocation away from the sex with a greater degree of competition among siblings (Lloyd, 1982; Clark, 1978; Bulmer and Taylor, 1980a; Frank, 1986). Accordingly, hermaphrodites in populations subject to local mate competition should bias allocation to their female function. Female-biased allocation is thus predicted for hermaphroditic plants that disperse pollen to a limited number of receptive flowers (Lloyd, 1982), or for plants in partially inbreeding populations (Charnov, 1982; Charlesworth and Charlesworth, 1981; Cutter, 2019; Sicard and Lenhard, 2011). The expected female-biased sex allocation of inbreeding hermaphrodites is well supported by empirical data, particularly comparisons between populations or species that have different rates of self-fertilization (West, 2009; Brunet, 1992; Schärer, 2009). Plant populations that have evolved high selfing rates often quickly evolve a ‘selfing syndrome’, including reduced allocation to pollen production and dispersal (Cutter, 2019; Sicard and Lenhard, 2011; Lemen, 1980). Such female-biased allocation in partially selfing plants exemplifies the outcome of selection under local mate competition (or ‘local sperm competition’; Schärer, 2009; Schärer and Pen, 2013; Yin and Haag, 2019; Cutter, 2015), where pollen grains (or sperm) from the same individual compete to fertilize a restricted pool of ovules (or eggs).

Self-fertilization represents the most extreme form of inbreeding, but inbreeding can also be the result of strong population structure. In demographically stable plant populations, mild inbreeding can occur if seeds and pollen are dispersed over short distances, because mating then occurs between individuals that are more closely related to one another than those drawn randomly from the population (Vekemans and Hardy, 2004). Stronger inbreeding may occur in species subject to metapopulation dynamics, i.e., frequent local extinctions and recolonizations (Whitlock and McCauley, 1990). This case is particularly interesting because individuals in a large randomly mating population may still be strongly inbred relative to the species or metapopulation as a whole, especially if the population was recently founded by one or a few individuals (Ives and Whitlock, 2002). This is because all individuals will be closely related through their descent from recent colonizers (Ives and Whitlock, 2002; Haag et al., 2002, 2005; Pannell and Obbard, 2003). In such situations, metapopulation dynamics may select for female-biased sex allocation at the species level, even if the mating system of local demes is fully outcrossing - mitigated to the extent that pollen dispersal among demes does not quickly erase the genetic effects of the colonization bottleneck (see Discussion). To our knowledge, this prediction has not yet been modeled for hermaphrodites, though Aviles (1993) used simulations to show that metapopulation dynamics favored a bias in the sex ratio of dioecious species.

The prediction that selection should favor female-biased sex allocation in metapopulations of species that outcross in their local demes also raises the question of how inbreeding should be measured to test it. Female-biased sex allocation due to inbreeding within populations is expected to be associated with elevated measures of local inbreeding, such as *F*_IS_ (Hamilton, 1967; West, 2009; Charnov, 1982), but *F*_IS_ does not capture the effects of inbreeding due to population genetic differentiation caused by population turnover. Rather, we should expect female-biased sex allocation to be associated with elevated *F*_ST_ in outcrossing species subject to inbreeding due to metapopulation dynamics. Indeed, both subdivision of the species into small demes (which causes inbreeding relative to the species as a whole) and the turnover of populations (in which enhanced inbreeding is caused by local population bottlenecks and the resulting strong within-deme co-ancestry) are well known to affect *F*_ST_ (Whit-lock and McCauley, 1990; Wade and McCauley, 1988; Pannell and Charlesworth, 1999, 2000; Rousset, 2004). *F*_ST_ captures both effects of inbreeding and genetic differentiation, but it has been criticized as a measure of the latter because of its sensitivity to the mutation rate and to levels of within-population variation, and alternative metrics have thus been proposed to replace it, notably *G*_ST_ (Hedrick, 2005) and Jost’s *D* (Jost, 2008; Jost et al., 2018). *G*_ST_ is effectively *F*_ST_ standardized by its maximum value for a given within-deme heterozygosity *H*_*S*_, whereas Jost’s *D* measures genetic differentiation calculated on the basis of the effective number of alleles within and among demes. Although the relative merits of *F*_ST_, Jost’s *D* and *G*_ST_ have been much debated (Whitlock, 2011; Leng and Zhang, 2011; Pannell and Fields, 2014; Alcala et al., 2014; Verity and Nichols, 2014; Jost et al., 2018; Wang, 2012; Alcala and Rosenberg, 2019), we expect *F*_ST_ to provide a better predictor of selection on a metapopulation’s sex allocation than Jost’s *D* and *G*_ST_ because of its connection to inbreeding (reviewed in Whitlock, 2011). To our knowledge, this expectation has yet not been evaluated.

Here, we derive predictions of the equilibrium sex allocation in an hermaphroditic metapopulation as a function of the rate of population turnover, the rate of dispersal among populations, and the selfing rate within demes. Our results indicate that population turnover should select for female-biased sex allocation as long as migration among populations is insufficient to erase the genetic signatures of inbreeding brought about by colonization. Using our analytical model and individual-based simulations, we also show that the inbreeding coefficients *F*_IS_, *F*_ST_ and *F*_IT_ are better predictors of the sex allocation in hermaphroditic metapopulations than the alternatives *G*_ST_ and Jost’s *D. F*_IS_ is a good predictor of sex allocation in partially selfing metapopulations, whereas *F*_ST_ is a better predictor in metapopulations of species that outcross locally.

## 2 Model and methods

### 2.1 Life-cycle

We consider a population of hermaphrodite individuals divided among an infinite number of demes, each carrying *n* individuals. The population goes through an annual life cycle as follows. Demes go extinct with a probability *e*. In non-extinct demes, each individual produces pollen and ovules according to the investment 0 ≤ *z* ≤ 1 of resources into female function (so that 1 *− z* is invested into male function). We assume that the total number of ovules produced in the deme is large. A proportion *m*_p_ of pollen produced in extant demes is dispersed to other demes, and the remaining pollen (proportion 1 *−m*_p_) is dispersed locally. Each ovule within a deme is fertilized by a pollen grain from the same individual with probability *α*, and by non-self pollen (either local or immigrant) with a probability 1 *−α*, based on a model of lottery competition. After mating, seeds are dispersed and all adults die. Seeds are dispersed such that, of the *n* individuals able to establish for the next generation, proportions *m*_s_ and 1 *−m*_s_ are represented by immigrant seeds and seeds produced locally, respectively.

Demes that have become extinct are assumed to be re-colonized according to either a ‘migrant-pool’ or a ‘propagule-pool’ model (Slatkin, 1977). In the former, 1 ≤ *k* ≤ *n* colonists are drawn randomly from across the entire metapopulation (Slatkin, 1977), whereas, in the latter, 1 ≤ *k* ≤ *n* colonists are drawn from single random deme. The migrant-pool model is assumed in our analytical treatment, whereas results under the migrant-pool model are compared with those for the propagule-pool model using individual based simulations. All *k* seeds then germinate and mate locally to bring the deme to its local carrying capacity *n*. This bout of reproduction immediately after colonization means that demes go through two rounds of reproduction in their first season, one to re-establish the deme at its carrying capacity, and then again as one of all demes reproducing in the metapopulation, as described above. This assumption is of course contrived, but it substantially simplifies analysis. However, our simulations (see below) indicate that this assumption has little effect on predictions. Note also that we assumed a fixed proportion of immigrant pollen grains (*m*_p_) and seeds (*m*_s_) for all non-extinct demes, regardless of their sex allocation, again for greater mathematical tractability. Here, too, our simulations show that this assumption had no major qualitative impact on our results.

### 2.2 Evolutionary analysis

We are interested in the evolution of the sex allocation, *z*, which is the proportion of reproductive resources allocated to female function. We assume that the value of *z* is determined at a single diploid locus evolving via rare mutations with weak additive effects. The direction of evolution is inferred from the selection gradient, *s*(*z*), which tells us whether selection favors an increase (when *s*(*z*) > 0) or decrease (when *s*(*z*) < 0) in the trait in a population monomorphic for *z*. Accordingly, the population will gradually converge towards a singular trait value, *z*^*^, which satisfies *s*(*z*^*^) *=* 0 and *s*^*′*^(*z*^*^) < 0, where the prime refers to differentiation with respect to *z*. Such a trait value is said to be convergence stable. We do not consider mathematically whether a population monomorphic for *z*^*^ is also uninvadable, but we later show by individual-based simulations that, for all parameters investigated, evolutionary convergent singular trait values are also uninvadable (i.e., that, once the population has converged for *z*^*^, the trait distribution remains unimodally distributed around *z*^*^).

#### 2.2.1 Selection gradient

As highlighted by much previous work (Frank, 1998; Rousset, 2004, for textbook treatments), the selection gradient in the infinite island model of dispersal in monoecious diploids under additive effects can readily be computed as

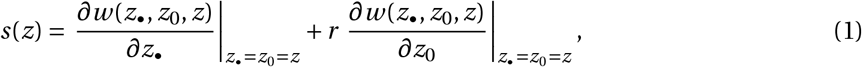

where *w* (*z*_*•*_, *z*_0_, *z*) is the fitness (i.e., the total expected number of successful offspring produced over one iteration of the life-cycle) of a focal individual with trait *z*_•_, when its *n* − 1 neighbors have on average trait *z*_0_, and the rest of the population is monomorphic for *z*, and *r* is a relatedness coefficient, measuring how likely a neighbor to a focal will transmit the same allele, relative to the focal individual itself transmitting this allele (in a monomorphic population). Specifically, *r* is defined here as

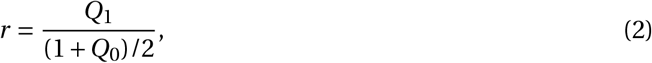

where *Q*_1_ is the probability that two neutral homologous genes in two different individuals from the same deme are identical-by-descent (IBD), while *Q*_0_ is the probability that the two neutral homologous alleles of one individual are IBD (which corresponds to the absolute coefficient of inbreeding under the infinite island model). The first term of eq. (1) corresponds to the effect of a change in sex allocation of the focal individual on its own fitness, i.e., the direct fitness effect of the trait, while the second term corresponds to the relatedness-weighted indirect fitness effect (i.e., the effect that a change in sex allocation in neighbors has on the fitness of a focal). To compute the selection gradient on sex-allocation, we specify below the fitness and relatedness for our life-cycle described above.

#### 2.2.2 Fitness

The fitness of an individual can be decomposed according to the number of offspring it produces via female function, *w*♀(*z*_•_, *z*_0_, *z*) (i.e., seeds that successfully establish), and via male function, *w*♂(*z*_•_, *z*_0_, *z*),

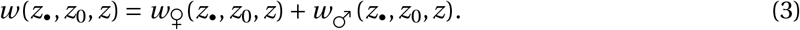

##### Fitness via female function

Under our assumption that all ovules are fertilized to produce seeds, fitness through female function can be written as the sum of seeds that successfully establish locally, in non-extinct demes, and in extinct demes, i.e.,

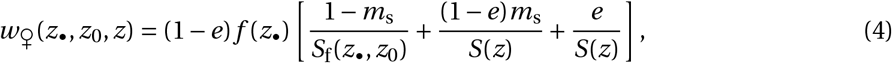

where (1 *−e*) is the probability that the focal deme survives, *f* (*z*_•_) is the number of seeds produced by an individual investing *z*_•_ resources into female function, which compete for a proportion (1 *−m*_s_) of spots in the focal deme, *m*_s_ in non-extinct demes (of which there are (1 *−e*)), and 1 in extinct demes.

The functions

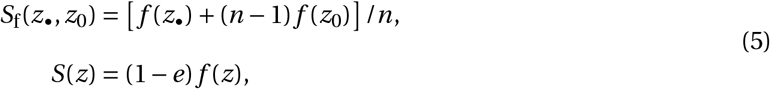

are the total number of seeds per individual that compete with the focal individual’s seeds in the focal deme and other demes, respectively.

##### Fitness via male function

The fitness of an individual through pollen is more complicated, as we need to take into account the ovules it fertilizes locally and those fertilized in other demes via pollen dispersal, as well as, in turn, whether the seeds it fertilizes establish or not, i.e.,

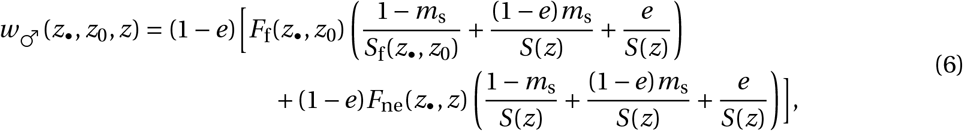

where *F*_f_(*z*_•_, *z*_0_) and *F*_ne_(*z*_•_, *z*) are the expected number of ovules available for fertilization by the pollen of a focal individual in the focal deme and in another (extant) deme, respectively. If *g* (*z*_•_) is the amount of pollen produced by an individual with sex allocation *z*_•_, these values can be written as

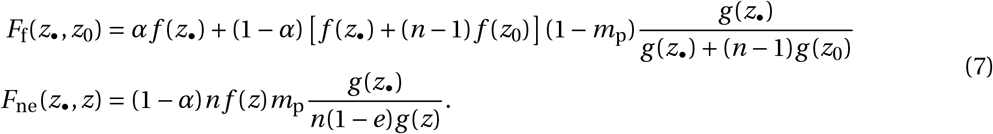

These expressions can be understood as follows. In its own deme (first line of eq. 7), an individual’s pollen fertilizes *αf* (*z*_•_) of its own ovules, and has access to (1 −*α*) *f* (*z*_•_) *+* (*n* − 1) *f* (*z*_0_) (1 −*m*_p_) other ovules, for which it competes with other local pollen (so that it fertilizes each of these ovules with a probability given by the fraction on the first line of eq. 7). In another extant deme (second line of eq. 7), meanwhile, an individual’s pollen has access to (1 − *α*)*n f* (*z*)*m*_p_ ovules, and fertilizes each of these ovules with a probability given by the fraction on the second line of eq. (7) (i.e., by the ratio of the focal individual’s pollen to the total amount of immigrating pollen in that deme). To focus on the effects of metapopulation dynamics, we assume a linear relationship between sex allocation and the quantity of ovules and pollen produced (i.e., *f* (*z*) ∝ *z* and *g* (*z*) ∝ 1−*z*). This generates useful insights into how sex allocation evolves in response to various demographic and ecological factors.

#### 2.2.3 Relatedness and *F* -statistics

The relevant probabilities of IBD under neutrality, *Q*_0_ and *Q*_1_, are computed by standard recursion methods (Karlin, 1968). We first define

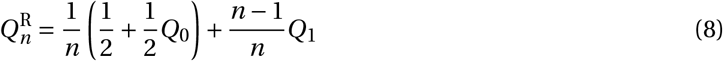

as the probability that two genes in two individuals sampled with replacement from a deme with *n* individuals are IBD. Then, at equilibrium, the probability *Q*_0_ that the two homologous alleles at a particular locus are IBD satisfies

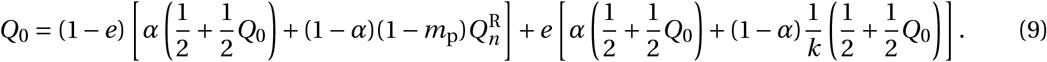

Eq. (9) can be read as follows. In an extant deme (probability 1 − *e*), two homologous alleles at a locus in a progeny derived from selfing (probability *α*) are IBD either if they are the same parental copy (with probability 1/2, assuming random segregation), or if the two parental alleles are themselves IBD (probability *Q*_0_/2). When the focal progeny is the product of random mating (with probability 1 −*α*), its two alleles are IBD only if the progeny’s paternally inherited allele is from the same deme (probability 1 −*m*_p_) and the two parental alleles are IBD (with probability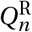). In a deme just recolonized after extinction (probability *e*), the two alleles in a selfed progeny are IBD with a probability also given by 1/2 +*Q*_0_/2. In a progeny produced by random mating, however, this probability of IBD depends on the number *k* colonizers that established the deme and reproduced locally, i.e., (1/2 +*Q*_0_/2)/*k*.

The probability *Q*_1_ that genes sampled from two individuals in the same deme are IBD can also be decomposed according to whether these individuals are sampled from a deme recolonized after extinction in the previous generation. Specifically, we have

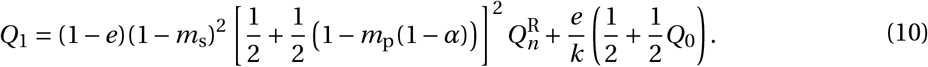

In an extant deme, two individuals carry IBD alleles only if both individuals are from the same deme (with probability (1 − *m*_s_)^2^). Then, with probability [1/2 + (1 −*m*_p_(1 −*α*)) /2] ^2^, two randomly chosen alleles from each of those two individuals were also present in adults within that same deme in the previous generation (for paternally inherited alleles, these alleles must not have immigrated into the deme). In turn, these two alleles are IBD with probability 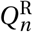. Finally, in a deme just recolonized after extinction, two individuals have IBD genes only if they share at least one parent (with probability 1/*k*), in which case two alleles sampled from both of these individuals are IBD with probability 1/2 +*Q*_0_/2.

Substituting eq. (8) into eqs. (9) and (10), and solving those jointly for *Q*_0_ and *Q*_1_ yields the equilibrium probabilities of IBD that are necessary to compute the coefficient of relatedness (eq. 2). This is easily achieved using an algebraic computer program but does not yield simple expressions in general. In addition to being necessary to compute the selection gradient, the probabilities *Q*_0_ and *Q*_1_ are closely connected to Wright’s *F* -statistics, which summarize the genetic structure of a subdivided population. Specifically, in the infinite island model of dispersal, *F*_ST_, *F*_IT_ and *F*_IS_ are respectively given by,

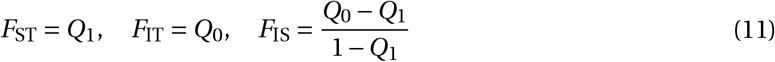

(Rousset, 2004, p. 18).

## 3 Results

### 3.1 Female-biased allocation favored by high selfing rate or small demes

In the absence of deme extinction (*e* = 0) and pollen dispersal (*d*_p_ = 0), we find that there exists a unique evolutionary convergent sex allocation,

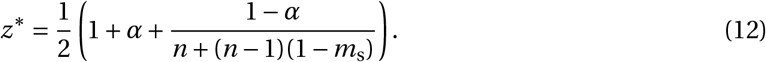

This strategy is increasingly biased towards female function (*z*^*^ > 1/2) as the rate of selfing *α* increases, especially when demes are large (i.e., *n* is large). This is because, under our assumption that all ovules are fertilized and there is no pollen limitation, investing into female function increases fitness via both male and female functions with selfing (*α* > 0). When demes are small, kin competition among pollen from the same parent to fertilize outcrossing ovules reduces selection for allocation towards male function (so that under random mating *α =* 0 and under complete seed dispersal *m*_s_ *=* 1, *z*^*^ *=* 1/2(1 + 1/*n*), which increases as *n* becomes small).

### 3.2 Extinction bottlenecks promote female-biased allocation

To investigate the effect of metapopulation dynamics on the evolution of sex allocation, we first consider a case in which there is no pollen dispersal among demes (*m*_p_ *=* 0), seed dispersal is weak and demes are large (specifically, *m*_s_ ∼ 𝒪 (*δ*) and *n* ∼ 𝒪 (1/*δ*), where *δ* is small parameter). Under these assumptions, the evolutionary convergent sex allocation is

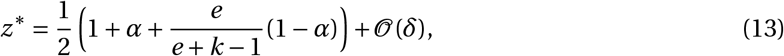

which increases as *e* increases and the number of colonizers *k* decreases. Frequent extinction and severe colonization bottlenecks thus favors the evolution of female-biased sex allocation because individuals producing more ovules and seeds are more likely to participate in the colonization of demes following their extinction. This effect is amplified when the number of colonizers *k* is low, e.g., when *k =* 1 a seed that successfully establishes a new deme will germinate, self and fill the whole deme with its own offspring.

Investigating *z*^*^ numerically confirms that extinction bottlenecks favor female-biased sex allocation, including when seed dispersal is not negligible (Fig. 1 where *m*_s_ is fixed to 0.1). Note that, with migrant-pool colonization (in which colonists are drawn independently from across the metapopulation), colonization by more than one (*k* > 1) individual contributes to gene flow among demes and thus reduces inbreeding due to population turnover and, in turn, selection for female-biased sex allocation. This dependence of inbreeding at the metapopulation level on the mode of colonization was first described by Slatkin (1977). Seen from the perspective of the sex allocation, when multiple colonizers from different origins establish a deme, the ovules of individuals that invest more into female function are more likely to be fertilized by unrelated competitors that invest more into male function, countering selection for female-biased sex allocation. This feature helps to explain why higher selfing rates, which prevent individuals from being fertilized by others, can lead to female-biased sex allocation even under weaker bottlenecks (*k* > 1, Fig. 1B). For comparison, we also modeled the case of propagule-pool colonization (where all *k* colonists hail from the same deme), and found that selection favors more extremely female-biased sex allocation, as expected (Fig. S3).

**Figure 1.**
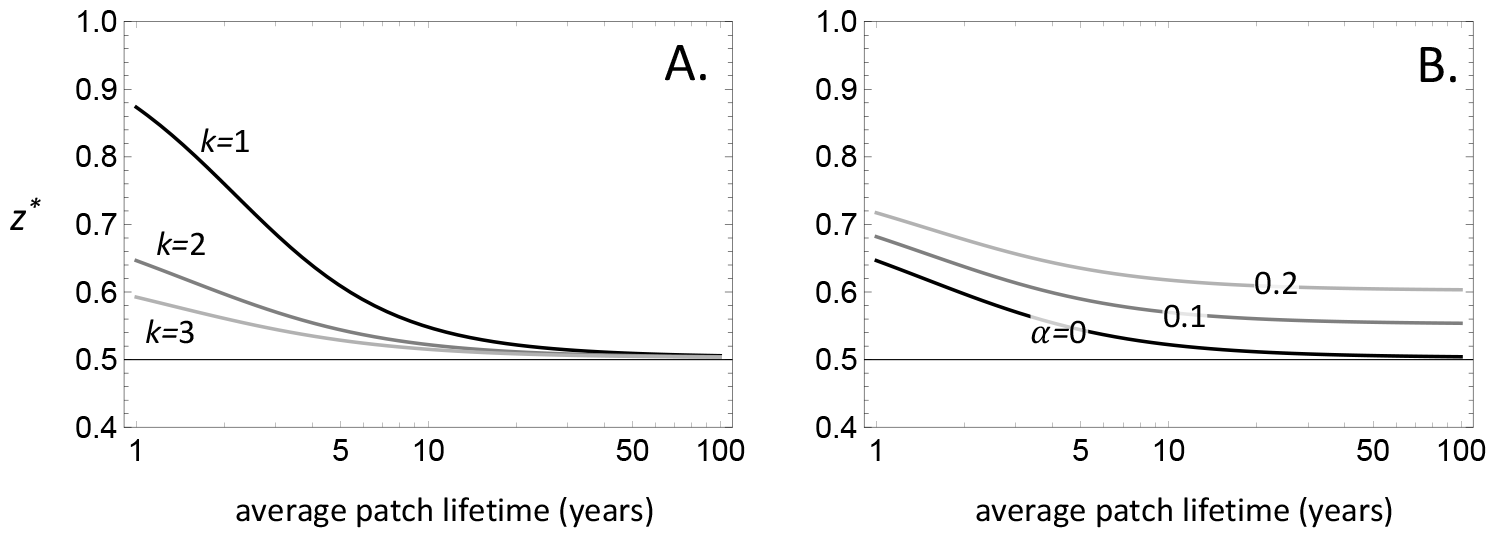
The effect of colonization bottlenecks and selfing on convergence stable sex allocation. Convergence stable sex allocation, *z*^*^, is plotted against the average deme lifetime, (1 − *e*)/*e*, in generations (shown on a log scale). The equilibrium strategy *z*^*^ is found by solving *s*(*z*^*^) = 0 for *z*^*^ from eq. (1). **A**. The effect of the number of colonizers, *k* = 1 (black), 2 (dark gray), 3 (lighter gray; other parameters, *n* = 100, *m*_s_ = 0.1, *m*_p_ = 0, *α* = 0). **B**. The effect of the selfing rate, *α* = 0 (black), 0.1 (dark gray), 0.2 (lighter gray) when *k* = 2 (other parameters, same as A).

### 3.3 Pollen dispersal among demes favors male-biased allocation

Next, we allow for pollen to disperse among extant demes. The expression for the convergence stable sex allocation under this scenario is too complicated to offer any analytical insights in the general case. However, when both seed and pollen dispersal are weak, selfing and extinction are rare, and demes are large (i.e., *m*_s_ ∼ 𝒪 (*δ*), *m*_p_ ∼ 𝒪 (*δ*), *α* ∼ 𝒪 (*δ*), *e* ∼ 𝒪 (*δ*) and *n* ∼ 𝒪 (1/*δ*) where *δ* is small parameter), the evolutionary equilibrium can be written as,

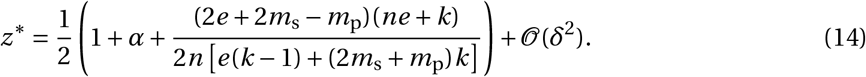

This equation reveals that, depending on the difference between pollen and seed dispersal, selection can favor either male-or female-biased sex allocation. Specifically, when *m*_p_ > 2(*e + m*_s_), selection promotes male-biased allocation, and female-biased allocation otherwise.

Computing the convergence stable sex allocation for various pollen and seed dispersal probabilities shows that the difference between those remains relevant when dispersal is not weak and extinctions are frequent, with greater seed dispersal favoring female-biased allocation (see red region in Fig. 2A). The relative value of seed versus pollen dispersal, however, is less relevant for selection on sex allocation when individuals self-fertilize their progeny, because selfing overwhelmingly favors female bias (Fig. 2B-C). In addition, pollen dispersal has little effect on the magnitude of the sex-allocation bias (i.e., the value of *z*^*^ rather than whether it is ≤ 1/2) unless extinctions are very frequent (Fig. 2D).

**Figure 2.**
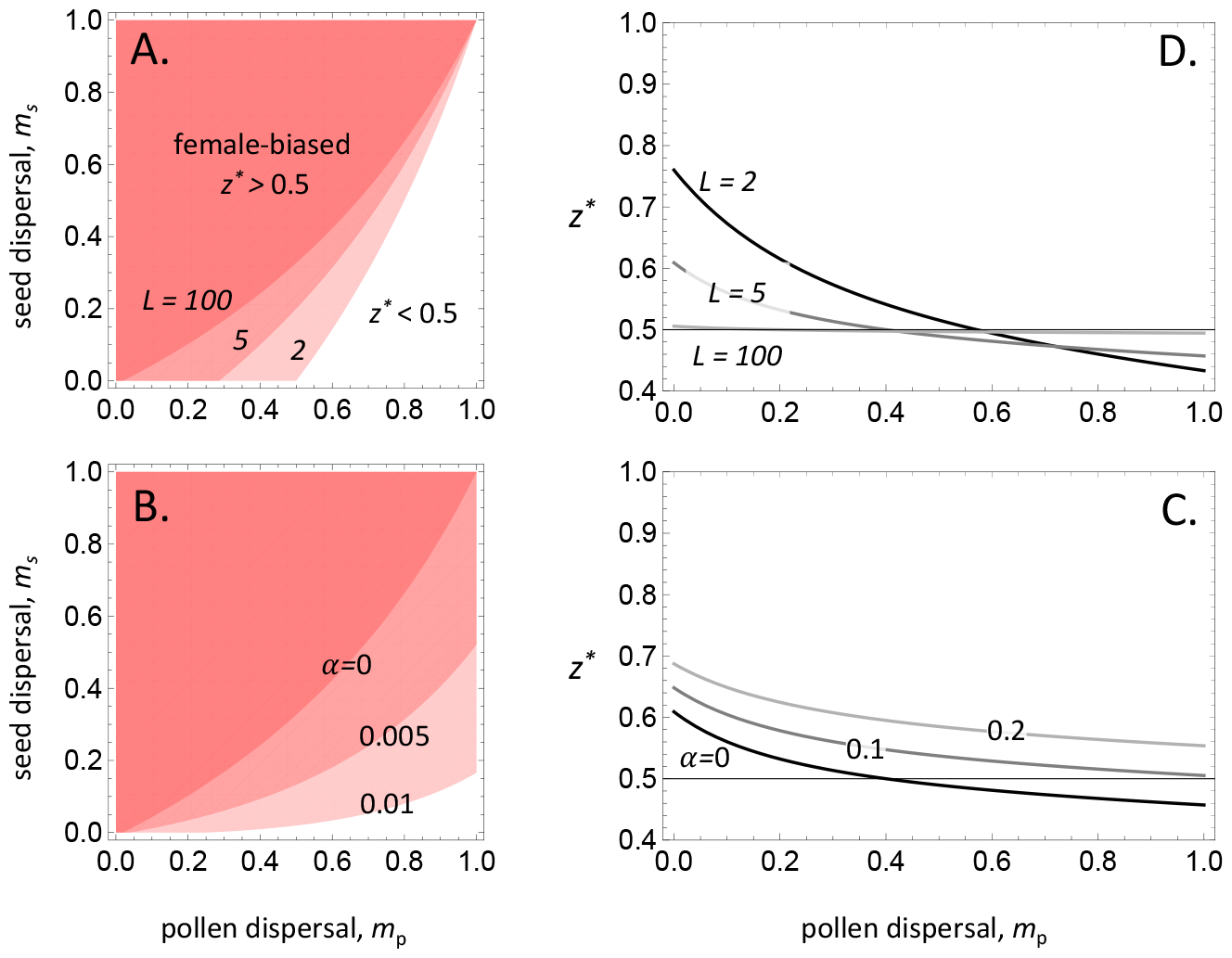
The effect of pollen dispersal on convergence stable sex allocation. **A.** Parameter region for which convergence stable sex allocation, *z*^*^, is biased towards female (*z*^*^ > 1/2, red region) or male function (*z*^*^ < 1/2). Different contours are shown for different average deme lifespans, *L* = 2, 5, 100 (other parameters: *n* = 100, *k* = 1, *α* = 0). **B**. Same as A but with different contours for different selfing rates, *α* = 0, 0.005, 0.01 (other parameters: *n* = 100, *k* = 1, *L* = 100). **C**. Convergence stable sex allocation, *z*^*^, for different selfing rates, *α* = 0 (black), 0.1 (dark gray), 0.2 (lighter gray; other parameters, *n* = 100, *m*_s_ = 0.1, *L* = 5, *k* = 1). **D**. Same as C for different average deme ages, *L* = 2 (black), 5 (dark gray), 100 (lighter gray) when *α* = 0 (other parameters, same as C).

### 3.4 Association between sex allocation and *F*-statistics

To assess the relationships between hierarchical measures inbreeding (*F*-statistics) and convergence stable sex allocation in the metapopulation, we generated random values for each model parameter over intervals that might reasonably occur in a natural metapopulation. We then plotted *F*_IS_, *F*_ST_ and *F*_IT_ against the sex allocation for three contrasting scenarios: the full model, including variation among simulations in the rates of selfing as well as pollen dispersal among demes; a model assuming random mating in local demes as well as pollen dispersal; and a model assuming random mating in demes but excluding pollen dispersal among demes (Fig. 3). The results indicate that when variation among metapopulation runs includes variation in the selfing rate of individuals, *F*_IS_ is most narrowly associated with sex allocation and the association with *F*_ST_ is relatively poor. In contrast, *F*_ST_ and sex allocation are more closely related in models in which the selfing rate is zero, especially when gene flow among demes takes place only via seeds and not pollen. Results of individual-based simulations of our model for the relationship between *F*_IS_, *F*_ST_ and *F*_IT_ and sex allocation were very similar (see Section A and Fig. S2 in the Supplementary Information). To compute Jost’s *D* and *G*_ST_, we performed the same calculations on the results of individual-based simulations of the same three models, for the same range of parameter combinations and for both migrant-pool (Fig. S6) and propagule-pool colonization (Fig. S7). These simulation results recapitulate calculations based on the analytic treatment for *F*_IS_, *F*_ST_ and *F*_IT_, and they also indicate that Jost’s *D* and *G*_ST_ are particularly poorly associated with sex allocation across the metapopulation.

**Figure 3.**
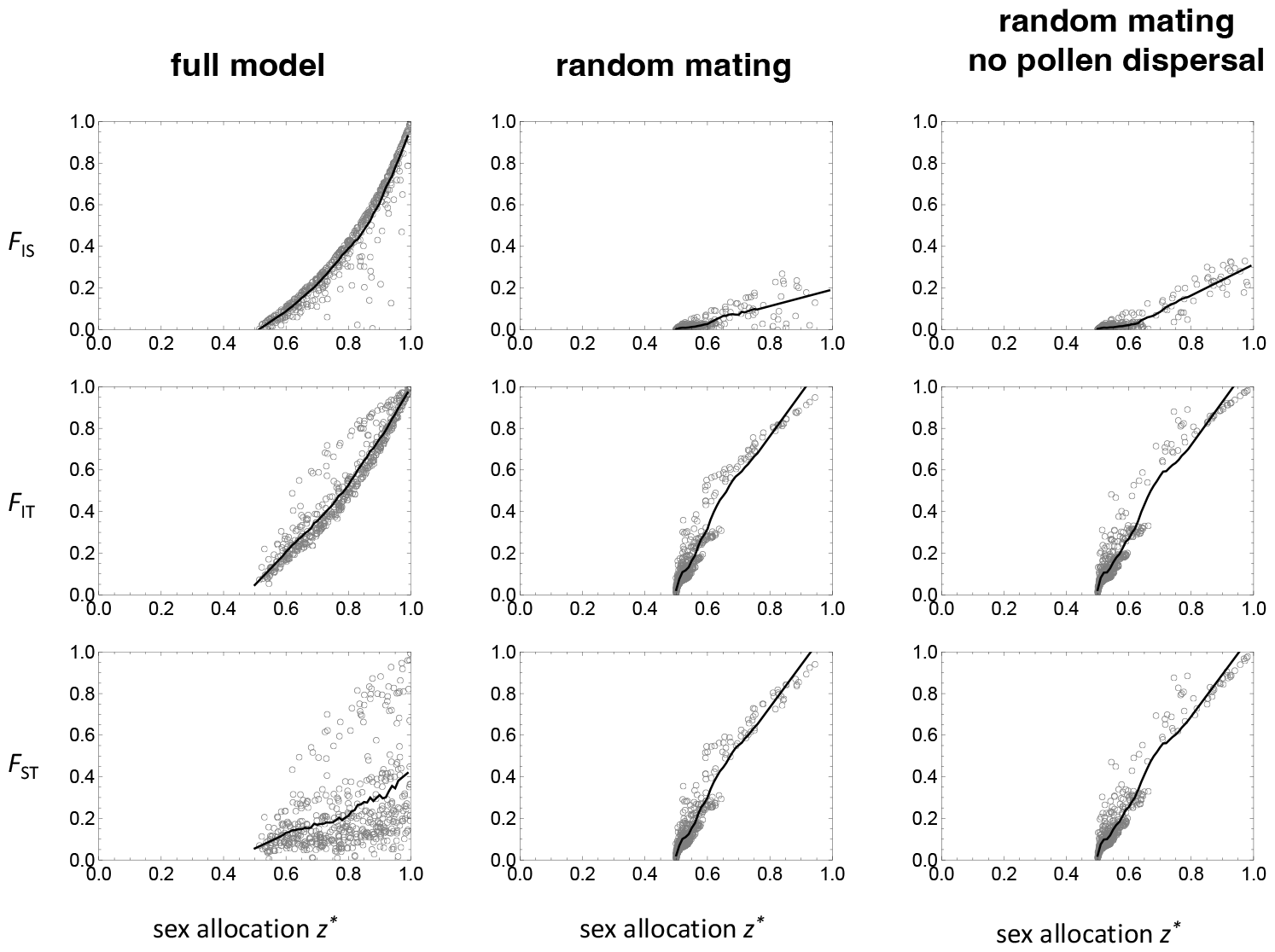
The association between sex allocation and *F*-statistics. Each grey empty circle gives the convergence stable sex allocation *z*^*^ and the associated *F*-statistic (first row *F*_IS_, second row *F*_IT_, third row *F*_ST_, calculated in adults, i.e., at the end or beginning of the life cycle) for a randomly generated set of parameters (with *n* = 1000 fixed). In black is the loess regression between *z*^*^ and *F*-statistic. First column: 500 sets of random parameter values with 1 ≤ *k* ≤ 6, 0 ≤ *α* ≤ 1, 0 ≤ *m*_s_, *m*_p_ < 0.1, and 0.005 < *e* < 0.5. Second column: 500 sets of random parameter values with same bounds as first column but with *α* = 0, i.e., under random mating. Third column: 500 sets of random parameter values with same bounds as first column but with *α* = 0 and *m*_p_ = 0, i.e., under random mating and in the absence of pollen dispersal.

### 3.5 Effect of competition between immigrants and local individuals on sex allocation

Our model (Section 2.1) assumes a fixed probability *m*_p_ that an outcrossed ovule is fertilized by immigrant pollen and a fixed probability *m*_s_ that a spot in an extant deme is filled by an immigrant seed. In other words, we assume the same level of competition between immigrants and local seeds or pollen grains. This rules out accounting for the possibility that individuals in a deme that on average invest more into male (or female) function receive proportionally less immigrant pollen (or seed) than those in other demes. In Section A of the Supplementary Information, we analyze a model in which each pollen grain and seed has a probability of emigrating away from its natal deme (*d*_p_ in pollen and *d*_s_ in seeds), so that demes that are more productive receive fewer immigrants. These two models of dispersal are typically referred to as backward (or effective) dispersal for *m*_p_ and *m*_s_, and as forward dispersal probabilities for *d*_p_ and *d*_s_. We find that the way selection shapes sex allocation differs little between these two models of dispersal for our assumed life-cycle (Fig. S1), leading to a similar association between sex allocation at evolutionary equilibrium and *F*-statistics (Fig. S2).

## 4 Discussion

### 4.1 Effects of population turnover on sex allocation in hermaphroditic metapopulations

Our study has demonstrated that population turnover in a metapopulation as a result of local extinctions and recolonizations, selects for female-biased sex allocation in hermaphrodites under a wide range of conditions that can be related to previous theory. The bias in allocation is strongest when the rate of turnover is high, new demes are colonized by small numbers of individuals from the same source population, population growth is rapid, and dispersal among extant demes is low. In contrast, the potential effects of population turnover on biased sex allocation are erased by migration, and when extinct demes are recolonized by individuals drawn from demes across the metapopulation. In general, these results are consistent with population genetic models that point to the importance of the relative rates of extinction and gene flow by migration for patterns of within-population diversity and among-population differentiation (reviewed in Pannell and Charlesworth, 2000). Our model indicates that this relativity extends to the evolution of sex allocation.

Our results align with well established theory on the effects on sex allocation of inbreeding in general, and of inbreeding due to population structure in particular (Hamilton, 1967). Biased sex ratios are predicted for and observed in dioecious parasitoids or fig wasps, in which a single or a few females lay their eggs in animal hosts or figs, respectively, and the hatching male progeny must compete among themselves to mate with their sisters (Werren, 1980; Frank, 1985; Herre, 1985, 1987). In such cases, over-investment in the production of males diminishes the inclusive fitness of an egg-laying mother, who would do better to use her limited eggs to produce more females. The idea that inbreeding should favor female-biased sex allocation in hermaphrodites is not new: local mate competition caused by self-fertilizing hermaphroditic plants, has long been interpreted as responsible for their low pollen-ovule ratios (Brunet, 1992; Charlesworth and Charlesworth, 1981; Lemen, 1980; Cruden, 1977; Lloyd, 1992), or their low allocation to pollinator attraction (Cutter, 2019), which may be viewed as benefiting male more than female function (Charlesworth and Charlesworth, 1987; Bell, 1985; Stanton et al., 1986; Barrett et al., 1996). Similarly, low inferred allocation to male function in hermaphroditic snails and worms has been attributed to local mate competition due to self-fertilization or biparental reproduction in genetically viscous populations (reviewed in Schärer, 2009; Yin and Haag, 2019). Our results build on these ideas by drawing attention to the capacity of metapopulation dynamics to bring about sufficiently strong inbreeding to bias a population’s sex allocation even in outcrossing hermaphrodite species.

The strong female biases our model predicts for hermaphroditic sex allocation in metapopulations are qualitatively similar to those predicted and reported for the sex ratios of dioecious invertebrates with strong population subdivision, strong local inbreeding, and frequent colony turnover (West, 2009; Hardy, 2002). In such species, the egg-laying mother is able to control the sex ratio of her progeny. In contrast, dioecious species with genetic determination of sex (i.e., in which sex is largely beyond maternal control) tend not to show the substantial biases in sex allocation observed in fig wasps, even though they may also be subject to demographic processes involving colonization and subsequent local inbreeding. An example would be a species with a lifestyle characterized by the ‘Haystack model’ first introduced by Maynard Smith (Smith, 1964). The Haystack model considers a population of mice that becomes periodically subdivided into small haystack habitat demes for several generations of mating among closely related co-habitants of the deme before haystacks are destroyed and the mice disperse to seek new demes; it is thus similar to the model we have simulated here. Bulmer and Taylor (1980a,b) and Taylor and Bulmer (1980) showed that female-biased sex ratios should evolve as a result of local mate competition in such populations. But mice, even if they were to live in a metapopulation of haystacks, seem less likely to realize the evolutionary stable sex ratios that theory would predict, because female mice probably have less control over the sex ratio than their fig-wasp counterparts (though see (Aviles et al., 2000) for an example of realized female bias in spiders with genetic sex determination). Similarly, although the ecology of the dioecious plants *Silene latifolia* (e.g., Moody-Weis et al., 2008), *S. dioica* (e.g., Giles and Goudet, 1997) and diploid *Mercurialis annua* (Dorken et al., 2017; Eppley and Pannell, 2007) involves population turnover and local inbreeding, sex ratios in these species are typically 1:1. We suggest that the processes we have modeled seem more likely to give rise to biased sex allocation in hermaphroditic species than in dioecious species in which sex is determined genetically. This is because selection adjusts the proportion of resources allocated to male versus female function by each individual, and a single monomorphic evolutionary stable sex allocation can be achieved.

Patterns of sex allocation displayed by monoecious and androdioecious populations of the annual colonizing plant *Mercurialis annua* are consistent with predictions of our model. *M. annua* is subject to population turnover (Dorken et al., 2017; Eppley and Pannell, 2007), sufficient to generate the strong population subdivision and inbreeding of the sort that should select for female-biased allocation (Obbard et al., 2006; Pannell et al., 2014). In the Iberian Peninsula, *M. annua* is a metapopulation comprising monoecious individuals that, in many populations, co-occur with males (Pannell, 1997b,a). Comparisons of the sex allocation of hermaphrodites with males reveal that the former are strongly female-biased (Pannell et al., 2014; Pannell, 1997a). Migration of males into populations of hermaphrodites lead to a rapid increase in their frequency, consistent with an initial deficit of male allocation at the population level (Dorken and Pannell, 2008). Importantly, hermaphrodites of *M. annua* are largely outcrossing (Korbecka et al., 2011), suggesting that their female-biased sex allocation is not an outcome of selection under habitual selfing but rather the effect of the metapopulation processes that typify this colonizing weed. It would be worth comparing the sex allocation of outcrossing hermaphrodites of other species among regions that differ in their history of colonization, e.g., between populations at the extreme edge of a range expansion and those at the core of the species’ range, or between the native and exotic ranges of invasive species.

### 4.2 Correspondence between sex allocation and measures of inbreeding and population differentiation

We also considered how the evolutionary stable sex allocation of a hermaphroditic metapopulation relates to measures of inbreeding and population differentiation. In general, previous theory and empirical work have established the dependence of the ESS sex allocation on within-population inbreeding (Hamilton, 1967; West, 2009; Charnov, 1982). For instance, the reduced allocation to male function in partially or wholly self-fertilizing species is understood to be the result of the local mate competition. Hermaphroditic plants with a colonizing habit, and thus subject to metapopulation dynamics, are often self-fertilizing and show allocation patterns consistent with female bias, particularly low pollen-ovule ratios (Sicard and Lenhard, 2011). Although we expect selection under self-fertilization to favor female-biased allocation, our simulations should caution us away from invoking within-population selfing as the sole basis of selection for such bias, because a history of population turnover may also have contributed to it.

A key insight from our study is that indices of inbreeding and population structure, such as *F*_IS_, *F*_IS_ and *F*_ST_ will differ in the extent to which they should predict sex-allocation bias on hermaphroditic metapopulations. In species that self-fertilize during the process of colonization but that outcross thereafter when mates become available with the growth of their population (as assumed in most of our simulations), *F*_IS_ will be low, reflecting the local outcrossing rate, despite the inbreeding brought about by colonization bottlenecks. Yet selection at the metapopulation level will favor female-biased sex allocation all the same, and this female bias should be associated with elevated *F*_ST_. Importantly, elevated *F*_ST_ provides a reasonable prediction of female-biased sex allocation irrespective of the underlying demographic causes, i.e., irrespective of whether *F*_ST_ is elevated because of high population turnover, low among-deme migration, or small numbers of colonists, or because colonists come from one rather than from many demes. This is useful because inferring the demographic processes on the basis of a genetic measure such as *F*_ST_ should normally be much easier than doing so on the basis of direct observation. For instance, our predictions of female-biased sex allocation due to metapopulation processes could be tested by comparing measures of *F*_ST_ among different geographic regions of a largely outcrossing species with an ability to self-fertilize under mate limitation: in regions subject to greater population turnover, e.g., at range margins, we would expect to see elevated *F*_ST_ and greater relative allocation to female function.

Our results emphasize the superiority of *F*_ST_ over other measures assessed for relating sex allocation to population structure, especially Jost’s *D*. The close relationship between sex allocation and *F*_ST_ illustrates its tight connection to evolutionary theory in general and to inbreeding in particular (Whitlock, 2011; Rousset, 2004; Taylor, 1988). Whereas *F*_ST_ measures deviations from random mating due to population structure, Jost’s *D*, which measures deviations from total genetic differentiation, is not connected in any formal way to a population’s history of inbreeding and is even rather insensitive to some processes that affect differentiation itself, such as genetic drift (Whitlock, 2011). Jost’s *D* is also slower to reach equilibrium than is *F*_ST_ (Ryman and Leimar, 2008), a fact likely to render it less useful for characterizing populations subject to demographic fluctuations in a metapopulation. Note too that whereas the dependence of *F*_ST_ on the migration rate is monotonically negative, irrespective of the extinction rate, the sensitivity of Jost’s *D* to the migration rate interacts in a complex way with the population turnover rate (a tendency that also applies to the normalized measures of *G*_ST_, though apparently less severely). The complex dependence of Jost’s *D* on the interaction between migration and extinction has not, to our knowledge, been noted previously, but it illustrates the inadequacy of Jost’s *D* to reflect the genetic outcome of evolutionary or demographic processes.

## Authors’ contributions

JRP conceived the study. CM developed the analytical model. CR wrote the simulation program and performed the simulations. JRP, CR and CM wrote the manuscript.

## Competing interests

The authors declare no conflict of interest.

## Acknowledgements

CR was funded by grants awarded to JRP by the grant Swiss National Science Foundation (31003A_163384) and by the University of Lausanne. CM was supported by Swiss National Science Foundation grant PCEFP3181243. We are grateful to Jérôme Goudet and Sam Neuenschwander for helpful discussions and implementation advice during the early stages of this project. The simulations were performed on the Core Cluster of the Institut Français de Bioinformatique (IFB) (ANR-11-INBS-0013).

## Supplementary Information

## A Forward dispersal model

Here, we analyse a model where each pollen grain and seed has a probability of emigrating away from its natal deme (*d*_p_ in pollen and *d*_s_ in seeds), so that, in contrast to our baseline model, demes that are more productive receive fewer immigrants.

### A.1 Set-up

#### A.1.1 Life-cycle

We change two steps of the baseline life-cycle elaborated in section 2.1. First, for step (3) pollen dispersal: each pollen grain either disperses to another randomly chosen deme (with probability *d*_p_) or remains in its deme of origin (with complementary probability 1 −*d*_p_). Second, for step (5) seed dispersal: each seed either disperses to another randomly chosen deme (with probability *d*_s_) or remains in its deme of origin (with complementary probability 1 −*d*_s_).

#### A.1.2 Fitness

These changes to the life-cycle require us to revisit the fitness functions used to derive the evolution of sex allocation (eqs. 4 & 6).

##### Fitness via female function

Under our new assumptions, fitness through female function is now written as

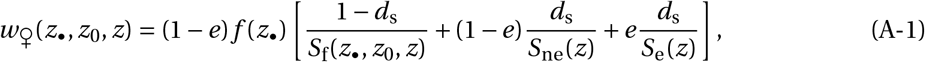

where (1 −*e*) is the probability that the focal individual’s deme survives, *f* (*z*_•_) is the number of seeds produced by an individual investing *z*_•_ resources into female function, (1 − *d*_s_) of which remain that in the focal individiual’s deme, (1 − *e*)*d*_s_ that disperse to a non-extinct deme and *ed*_s_ to an extinct deme. The functions *S*_f_(*z*_•_, *z*_0_, *z*), *S*_ne_(*z*), and *S*_e_(*z*) are the total number of seeds per individual before competition in the focal deme, in a non-extinct deme and in an extinct deme, respectively. These densities are given by

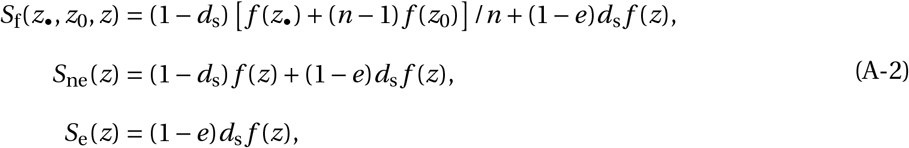

i.e., the sum of seeds that remain (except for extinct demes) or disperse from other demes.

##### Fitness via male function

The fitness of an individual through pollen, meanwhile, now reads as

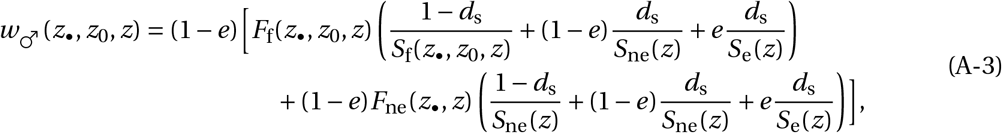

where *F*_f_(*z*_•_, *z*_0_, *z*) and *F*_ne_(*z*_•_, *z*) are the expected number of ovules fertilized by the pollen of a focal individual in its own deme and in another (non-extinct) deme, respectively. With *g* (*z*_•_) as the amount of pollen produced by an individual with sex allocation *z*_•_, these are

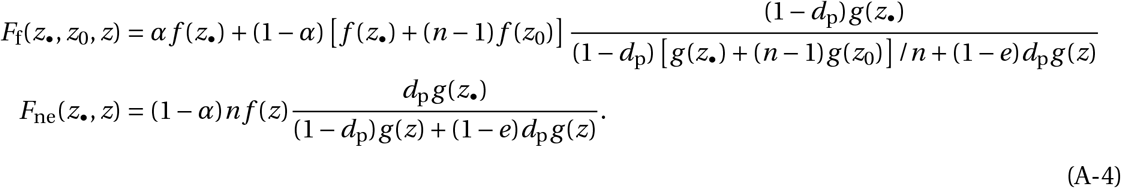

#### A.1.3 Relatedness

Our new assumptions about dispersal do not change the way we calculate relatedness (section 2.2.3). They require us, however, to express *m*_p_ and *m*_s_, which in eqs. (8) & (9) correspond to the probabilities that a successful pollen grain and seed come from another deme under neutrality, in terms of the forward probabilities of dispersal *d*_p_ and *d*_s_. These are given by

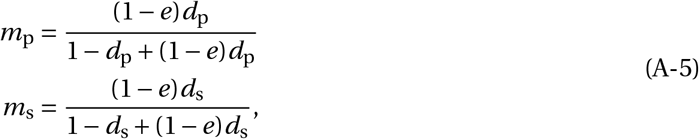

i.e., by the ratio of immigrants from non-extinct demes to the total number of dispersing units (pollen grain or seeds) in a non-extinct deme before competition (for fertilization or establishment).

### A.2 Results

The evolutionary convergent sex allocation is derived in the same way as for our model in the main text. Analysis of the solution reveals that considering forward dispersal does not change our main qualitative results. In particular, local bottlenecks lead to similar levels of female bias in sex allocation under both models of dispersal (compare figs. 1A and S1A). More generally, we observe very similar effects of seed and pollen dispersal on the sex allocation bias favored by selection, whether it is backward or forward (compare figs. 2A and S1B). In line with this result, the association between sex allocation and *F*-statistics under forward dispersal is almost indistinguishable from the case where backward dispersal is fixed (compare red and black dashed line in fig. S2).

## B Individual-based simulations

We used individual-based simulations to examine the evolution of a metapopulation under conditions identical to those assumed in Section 2.1, with an additional variable accounting for finite population growth (rather than growth from *k* colonizers to carrying capacity). In addition to considering a more realistic demography, these simulations also allowed us to consider compute Jost’s *D* and *G*_ST_ and thus to compare the association between these statistics and sex allocation with Wrights *F* statistics.

### B.1 Implementation

The model described in Section 2.1 was implemented in a forward individual-based simulator coded in C (link to GitHub repository: https://github.com/popgenomics/simulator_metapop_hermaphrodites#simulator-metapop-hermaphrodites).

#### B.1.1 Initialization of the metapopulation

The first generation starts with a metapopulation of already pre-established adults. This metapopulation is made up of *N*_d_ demes, all initially filled to carrying capacity *K* . Each of the initialized hermaphroditic individuals has a female allocation varying between 0 and 1, which is determined by the sum of the effects of the two alleles at a single locus. At the stage of metapopulation initialization, a value for each allele of each individual is randomly drawn from a Uniform distribution [0, 0.5], generating a first parent generation with various allocation values *z* in [0, 1]. At the same stage, neutral alleles are also randomly assigned to a number *N*_l_ = 20 of neutral loci, encoded by a positive integer in [0, 999].

#### B.1.2 Mating

Prior to reproduction, each deme is assigned an extinct or extant status, with probabilities *e* or 1 -*e*. Extinct demes do not contribute to pollen and seed dispersal. Every generation, reproduction occurs, deme by deme, according to the following steps:

1. The number of seeds *N*_s_ to be produced is determined for each deme. This number of local seeds is the sum of the seeds produced individually by each individual (see point below). If this sum over all individuals is greater than carrying capacity *K*, then the number of local seeds is equal to *K* (i.e., *N*_s_ *= K*).
2. Individually, the number of seeds produced is the product of the female allocation *z* and a fecundity parameter. For a fecundity of 300, say, individuals with a female allocation will produce 300 seeds for *z* = 1 but 150 seeds for *z* = 0.5. If the expected number of seeds produced is a decimal, then this number is rounded to the smallest integer with a probability of adding a seed to this number equal to 1 minus the fractional part of the expected number of seeds. For example, if *z* = 0.501 and fecundity = 300, then the expected number of seeds is 150.3, i.e., 70% chance of producing 150 seeds and 30% chance of producing 151 seeds.
3. For each deme, future mothers are drawn with replacement *N*_s_ times, with their probability of being chosen weighted by their female allocation *z*.
4. Future fathers are drawn with replacement from the whole metapopulation, with an individual’s weight depending on both the male allocation (1 − *z*) and whether the individual is in or outside the focal deme. Males from the focal deme have a weight of (1 − *z*)(1 −*d*_p_), while males from other demes have a weight of 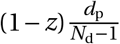, where *d*_p_ is the pollen dispersal rate and *d* is the number of demes in the metapopulation. A proportion *α* of seeds randomly drawn from the population are produced after self-fertilization. For these seeds, the father is the same individual as the mother.
5. Seeds are produced by drawing of one of two alleles for each neutral gene and each sex-allocation locus with equal probability. All loci are assumed to be genetically independent.

#### B.1.3 Seed dispersal

Only extant demes contribute to and benefit from seed dispersal. They receive a number of immigrant seeds drawn from a Poisson distribution with parameter *N*_imm_. For a given deme, seed migration follows a migrant-pool model, where each seed comes from an independent deme. Thus, for each immigrant seed, the original deme is drawn randomly from the population with a weight per deme corresponding to the number of produced seeds. To avoid slowing down the calculation time, the migrating seed is assumed to be a clone of a seed produced at the step B.1.2 in the donor deme, drawn uniformly from its cohort. Immigrant seeds that take the deme over its carrying capacity *K* randomly replace local individuals.

#### B.1.4 Recolonization

Demes that have become extinct are filled by *k* colonizers following either a propagule-pool or migrant-pool model.

1. Under propagule-pool colonization, the extinct deme is recolonized by *k* seeds from a single extant deme, randomly drawn from demes across the metapopulation weighted by the number of seeds they produce.
2. Under migrant-pool colonization, *k* independent draws of demes are made with replacement, weighted by the number of seeds they produce. Each sampled deme contributes one seed each time it is chosen.

Again, colonizers are assumed to be clones of newly produced individuals, as described for immigrants above.

#### B.1.5 Mutation and flowering

Each generation ends with the addition of mutations to all individual progeny seeds at different rates *μ*_n_ and *μ*_q_. For both categories of loci, mutations affect the maternal and paternal alleles with the same probability.

1. An allele at a neutral locus has a probability *μ*_n_ of mutating (based on binomial sampling). The mutation is implemented by drawing a new allele from a Uniform distribution in [0, 999].
2. Alleles at loci controlling the sex allocation *z* mutate at a rate *μ*_q_ (again, on the basis of a binomial draw). When this occurs, the current allelic value at the mutated locus is multiplied by a factor randomly drawn from a uniform distribution on [0.9, 1.1] until a limit value for the allocation of 0 or 1 is reached.

At the end of each generation, the parental generation dies out and seeds in the deme become adults with an allocation *z* depending on the post-mutation allelic values. Demes that become extinct in this generation are drawn by random sampling with probability *e*.

#### B.1.6 Returned values

For each run with a new combination of parameter values, the simulator returns the female allocation (mean and standard deviation) as well as the mean and standard deviation of 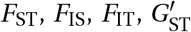 and Jost’s *D* (but see Section B.3).

### B.2 Multilocus estimators of inbreeding and population differentiation

For each neutral locus *i ∈* {1,…, *N*_l_}, we quantified from the simulations three probabilities of identity in state: *Q*_1,*i*_ (proportion of observed homozygous individuals in the whole metapopulation), *Q*_2,*i*_ (proportion of homozygous within demes, measured as its average over demes and weighted by the number of pairs of gametes within each deme), and *Q*_3,*i*_ (proportion of expected homozygotes in the whole metapopulation). Following Cockerham and Weir (1987), these are given by:

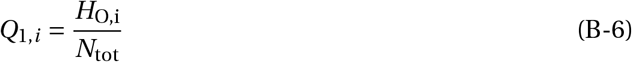

where *H*_O,i_ is the total number of homozygous individuals in the whole metapopulation at locus *i* and *N*_tot_ is the total metapopulation size (i.e. the number of diploid individuals);

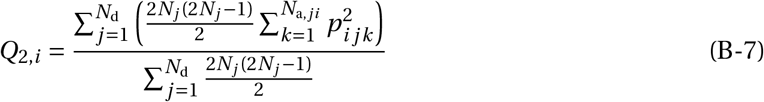

where *N*_d_ is the total number of demes, *N*_*j*_ is the number of individuals in deme *j, N*_a,*ji*_ is the number of alleles at locus *i* in deme *j*, and *p*_*i jk*_ is the frequency of allele *k* in deme *j* at locus *i*, 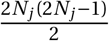 is a weighting factor approximated given by the number of gamete pairs in deme *j* assuming random mating within the deme; and

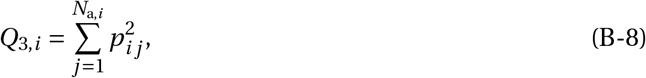

where *N*_a,i_ is number of different alleles at locus *i* in the metapopulation, and *p*_*i j*_ is the frequency of allele *j* at locus *i* in the metapopulation.

From these quantities measured individually for each of the *N*_l_ neutral locus, we obtain the global multilocus estimators of *F*_ST_, *F*_IS_ and *F*_IT_ following Weir (1996) and the implementation by Rousset (2008):

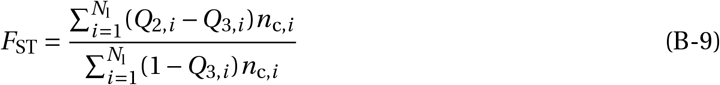

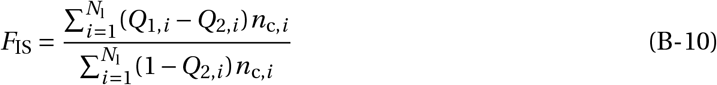

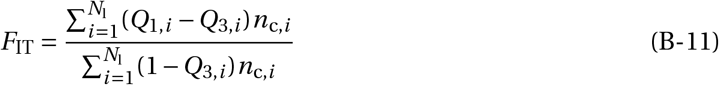

Where

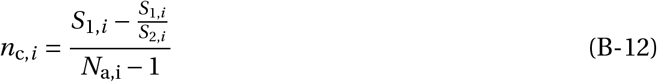

is a correction factor for the sample size at locus *i* that depends on *N*_a,i_ (which recall is the total number of alleles segregating in the whole metapopulation at locus *i*), and

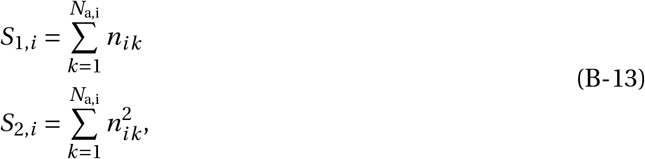

with *n*_*ik*_ the number of copies of allele *k* at locus *i* in the metapopulation.

In addition, we computed Jost’s *D*; and 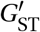 with a correction for diversity at individual loci as

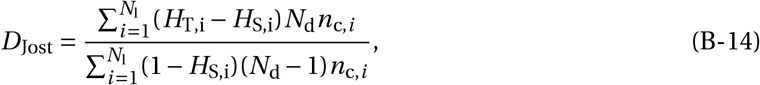

and

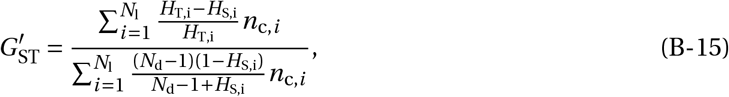

where *H*_S,i_ (the average expected heterozygosity across all demes) and*H*_T,i_ (the expected heterozygosity in the whole metapopulation) are measured at locus *i*, and *N*_d_ is the total number of demes.

### B.3 Association between sex allocation and measures of inbreeding and population differentiation

We used individual based simulations to compute Jost’s *D* and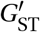, as well as F-statistics; simulation results were computed for scenarios of both migrant-pool (Fig. S6) and propagule-pool colonization (Fig. S7). The simulation results recapitulate calculations based on the analytical treatment (shown in Fig. 3), and also indicate that Jost’s *D* and 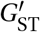 are particularly poorly associated with sex allocation across the metapopulation. To quantify our comparisons, we performed partial least squared regression (PLS; Abdi, 2010) to identify which statistics best explain the variance for *z* across simulations. Results are shown in Table 1 in the main text. The fitted PLS regression model explains between 94.7% and 97.1% of the total variance for *z* over the six sub-models explored. The demographic processes captured by the statistics calculated at neutral markers thus allow for robust predictions of the value for *z* at equilibrium. The latent components constructed with *F*_ST_, *F*_IS_, *F*_IT_, Jost’s *D* and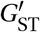, predict *z* accurately, with root mean squared error values ranging from 0.019 to 0.057. Finally, we assessed the importance of the ‘variables in the projection’ (VIP; Mehmood et al., 2020) of *z* for each of the six submodels. In models including selfing (specifically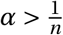), variation in *F*_IS_ is more narrowly associated with variation in *z* with VIP values (1.595 and 1.586 for models *i* and *ii* in Table 1) than *F*_ST_ (0.26 and 0.366 respectively), as for the analytic calculations. However, when the amount of self-pollen contributing to reproduction decreases (by decreasing *α* or increasing *m*_p_), *F*_IS_ becomes a relatively weak predictor (VIP values for *F*_IS_ equal to 0.519, 0.713, 0.587 and 1.097) compared to *F*_ST_ (1.41, 1.319, 1.92 and 1.277 respectively). 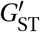 and Jost’s *D* are poorly associated with variation in female allocation.

**Table 1:**
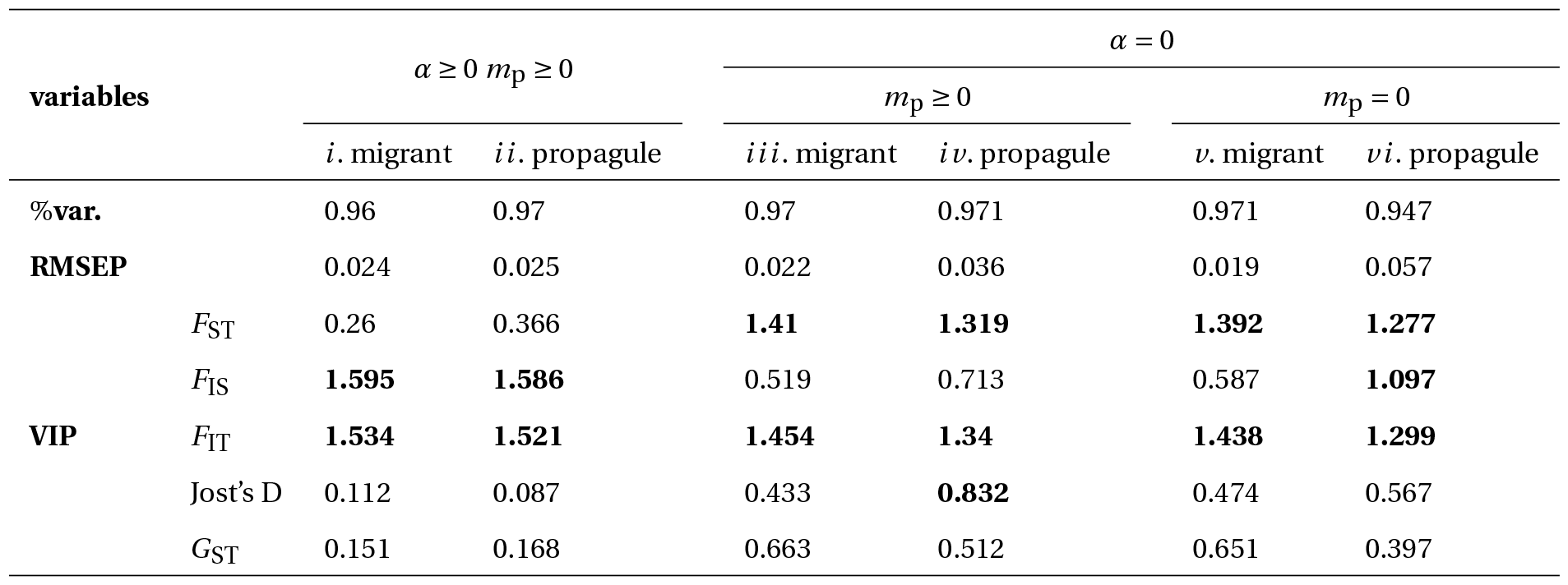
Strength of various statistics for inbreeding or population differentiation to predict the female sex allocation *z*. See Appendix B.3 for details on these analyses. “%var.” (first row) gives the percentage of variance in *z* explained by the latent components built on *F*_ST_, *F*_IS_, *F*_IT_, Jost’s *D* and *G*_ST_. RMSEP refers to the estimated root mean squared error of prediction of the fitted partial least squares (PLS) regression model. VIP refers to the variable importance in projection, based here on the weighted sums of the absolute regression coefficients. A PLS regression was performed for each of the six sets of simulations. For each set, 1,500 iterations were performed of a metapopulation of 1,000 demes of 1,000 individuals over 10,000 generations. Extinction *e* was sampled from a Uniform distribution *U* (0, 0.5), *k* from *R*(1, 6), *α* from *U* (0, 1), *m*_s_ and *m*_p_ from *U* (0, 0.1). The selfing rate *α* was set to zero for datasets from *i i i* to *vi* . Pollen dispersal *m*_p_ was set to zero for datasets *v* and *vi* . The terms migrant and propagule refer to the colonization models assumed.

### B.4 Effects of the slower growth of demes to their carrying capacity

We also used our individual based simulations to relax the assumption of our analytical model that colonizers of empty patches produce a large number of seeds so that demes are immediately taken to carrying capacity. This may be realistic for r-selected colonizers of small patches, but in general the growth of new demes to their carrying capacity may take several generations during which new immigrants may continue to arrive (thereby reducing the strength of the colonising bottleneck). To explore the effect of slower population growth on the evolution of sex allocation, we performed individual-based simulations under high or low fecundity (specifically, simulations where the maximum number of seeds produced is 300 and simulations where this is 3). Results are presented in Figure S4-A) for demes of size 100.

Low fecundity reduces the effects of bottleneck severity during recolonization on allocation bias only when the turnover rate is high (average deme lifetime = 1), but has little effect on the value of the trait when extinction is infrequent (average deme lifetime = 100). The results are qualitatively similar for migrant-pool and propagule-pool colonization, but with a consistently more female-biased allocation at equilibrium for propagule-pool colonization (Fig. S3-A), for all the explored values of selfing *α* (Fig. S5-B and S3-B). Similarly, a lower fecundity reduces the negative effects of pollen dispersal on female allocation, even when extinctions are frequent (Fig. S4-A). Finally, reduced fecundity leads to an equilibrium female bias close to that expected for a single deme in response to selfing alone, and this for any value of pollen dispersion (Fig. S4-B).

## Supplementary Figures

**Figure S1:**
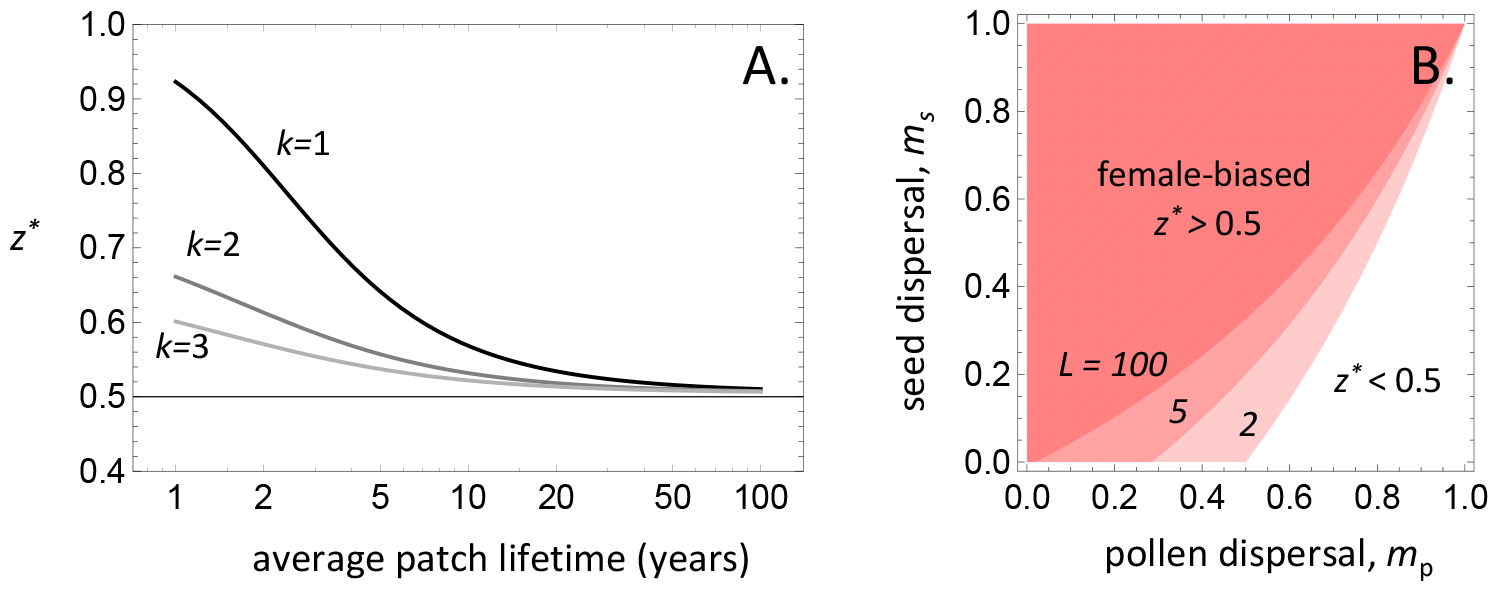
Convergence stable sex allocation with forward dispersal. **A**. Convergence stable sex allocation, *z*^*^, against the average deme lifetime, (1 −*e*)/*e*, in generations (shown on a log scale). The equilibrium strategy *z*^*^ was found by solving numerically *s*(*z*^*^) = 0 for *z*^*^ from eq. (1) with *k* = 1 (black), 2 (dark gray), 3 (lighter gray; other parameters, *n* = 100, *d*_s_ = 0.1, *d*_p_ = 0, *α* = 0). **B**. Parameter region for which the convergence stable sex allocation, *z*^*^, is biased towards female (*z*^*^ > 1/2, red region) or male function (*z*^*^ < 1/2). Different contours are for different average deme lifetimes, *L* = 2, 5, 100 (other parameters: *n* = 100, *k* = 1, *α* = 0).

**Figure S2:**
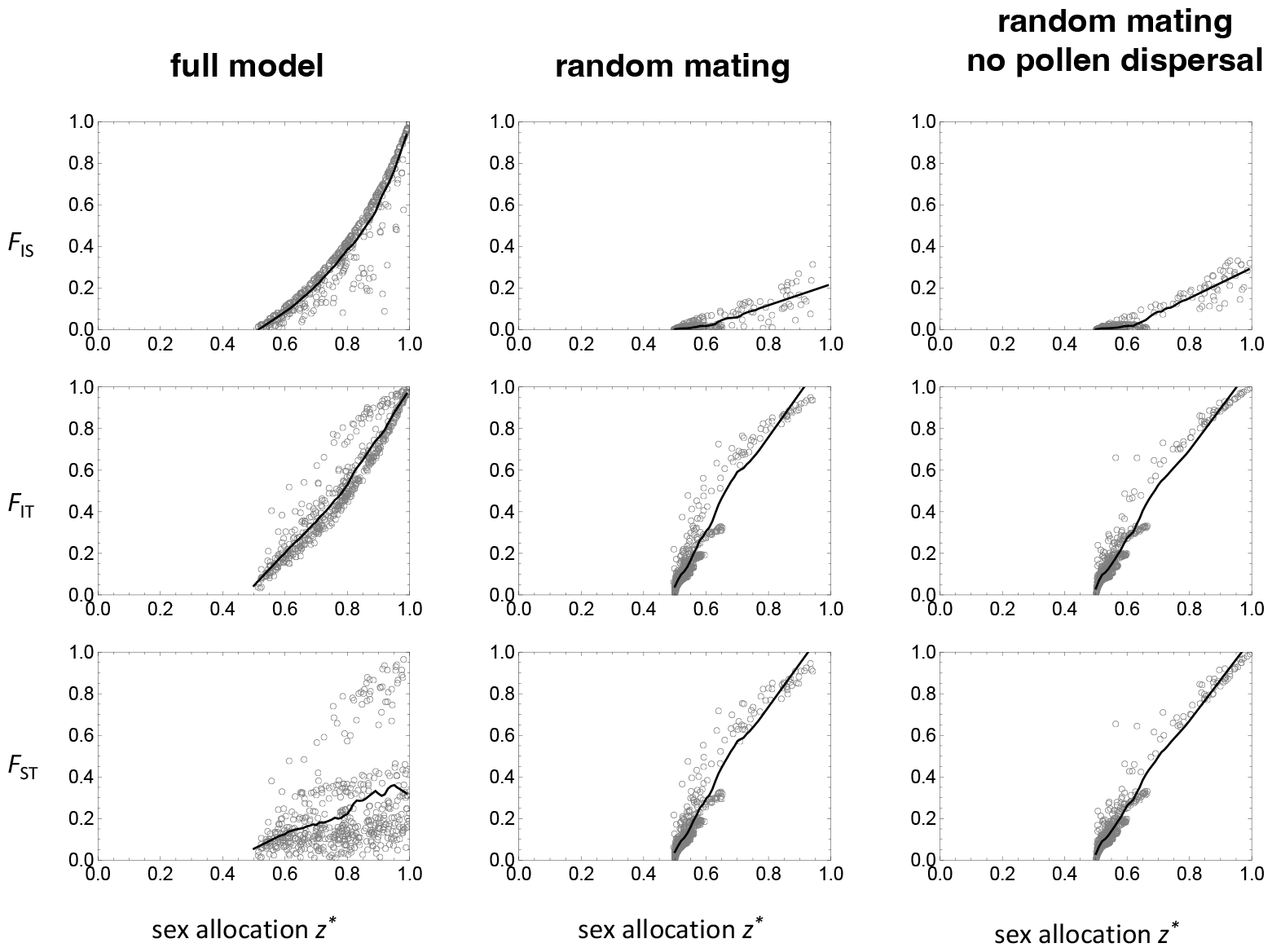
The association between sex allocation and *F*-statistics under forward dispersal. Each grey empty circle gives the convergence stable sex allocation *z*^*^ and the associated *F*-statistic (first row *F*_IS_, second row *F*_IT_, third row *F*_ST_, calculated in adults, i.e., at the end or beginning of the life cycle) for a randomly generated set of parameters (with *n* = 1000). In black is the loess regression between *z*^*^ and *F*-statistic. First column: 500 sets of random parameter values with 1 *≤ k ≤* 6, 0 *≤ α ≤* 1, 0 *≤ m*_s_, *m*_p_ < 0.1, and 0.005 *< e <* 0.5. Second column: 500 sets of random parameter values with same bounds as first column but with *α* = 0, i.e., under random mating. Third column: 500 sets of random parameter values with same bounds as first column but with *α* = 0 and *m*_p_ = 0, i.e., under random mating and absence of pollen dispersal.

**Figure S3:**
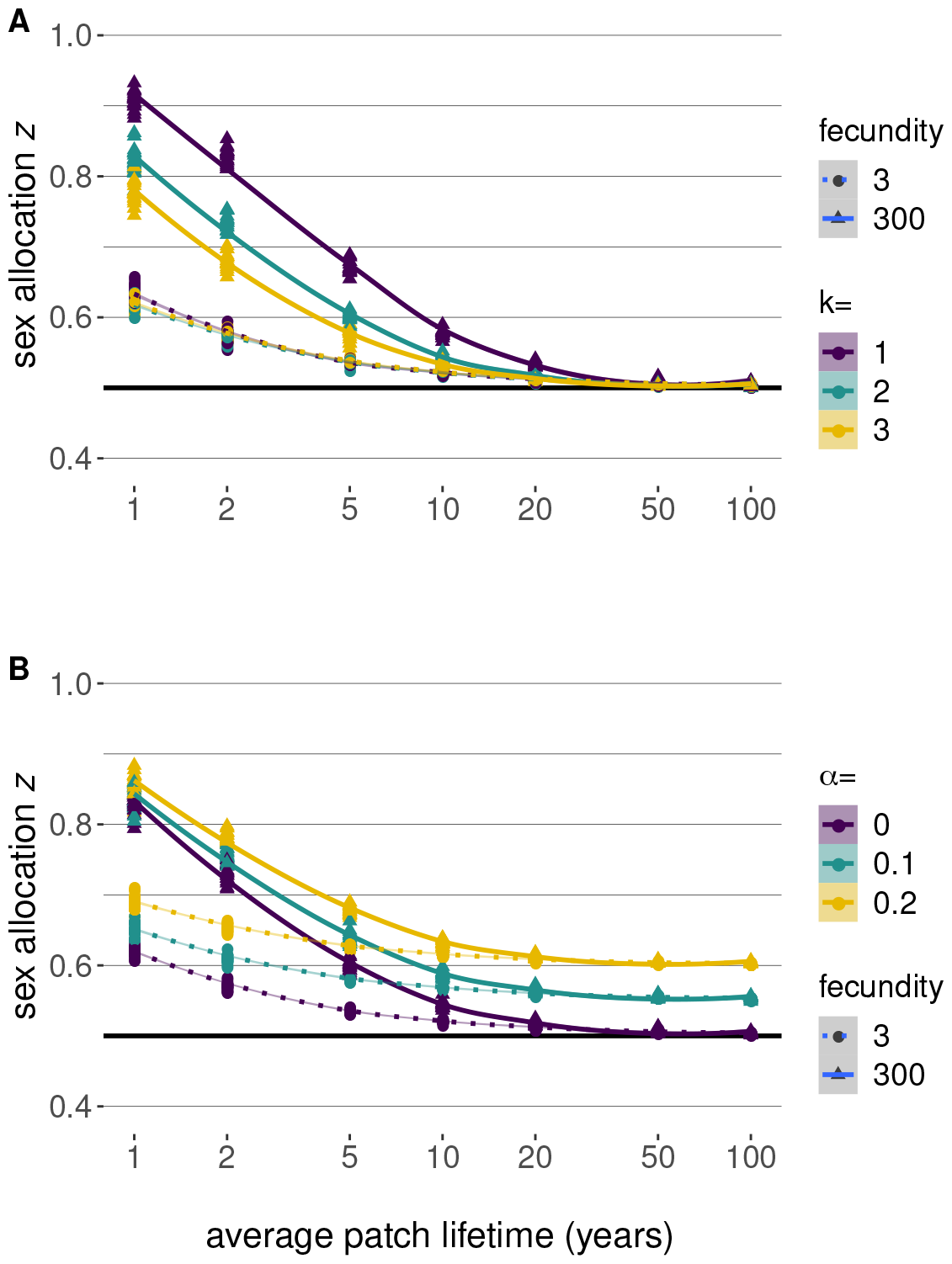
Simulations of the effect of extinction bottlenecks and selfing on convergence stable sex allocation in a model with propagule-pool colonization. A metapopulation of 1,000 demes of 100 individuals was simulated over 5,000 generations. 20 replicates were simulated for each combination of parameters. Contrary to analytical expectations (Figure 1), the extinct demes are here not immediately filled to carrying capacity by a specific round of reproduction of the *k* recolonising individuals. Fecundity is the maximum number of seeds a hermaphrodite can produce, i.e., when the female allocation = 1. Two values of fecundity were explored: fecundity = 3 (circles for individual simulations, dotted lines for the loess regression); and fecundity = 300 (triangles for individual simulations, solid lines for the loess regression). X-axis: extinction rate represented by the average lifetime of a deme. Y-axis: average female allocation in the metapopulation. Seed dispersal *d*_s_ = 0.05, *d*_p_ = 0 and selfing rate *α* = 0. **A**. seed dispersal *d*_s_ = 0.05, pollen dispersal *d*_p_ = 0, and selfing rate *α* = 0. After extinction, a deme could be recolonized by *k* = 1 (purple), *k* = 2 (blue) or *k* = 3 (yellow) individuals. **B**. dispersal similar to panel A, but with *α* = 0% (purple), *α* = 10% (blue) and *α* = 20% (yellow). After extinction, demes were recolonized by *k* = 2 individuals.

**Figure S4:**
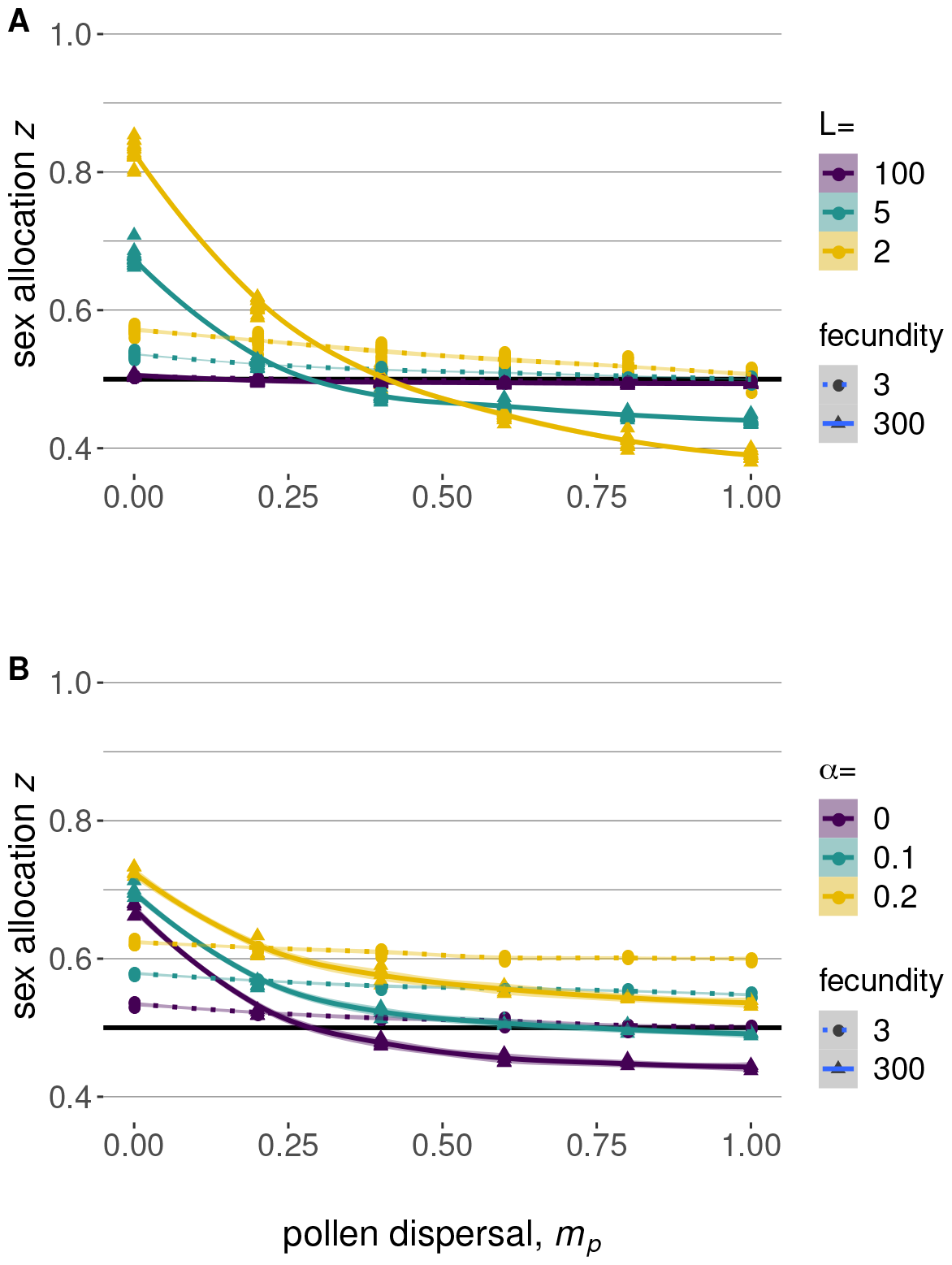
Effect of the degree of interdeme pollen dispersal on the female sex allocation in a metapopulation. A metapopulation of 1,000 demes of 100 individuals was simulated over 5,000 generations. 20 replicates were simulated for each combination of parameters. In contrast with analytical calculations for Figure 2, extinct demes are not immediately filled up to carrying capacity by a single round of reproduction of the *k* = 1 recolonizing individuals. Pollen dispersal refers to the fraction of outcrossed ovules fertilized by pollen coming from a different deme. **A**. Three extinction rates were explored for *α* = 0: average deme lifetime of 100 generations (purple); 5 generations (blue); 2 generations (yellow). **B**. Three selfing rates were explored for a deme life span fixed to 5 generations: *α* = 0 (purple), *α* = 10 (blue) and *α* = 20 (yellow).

**Figure S5:**
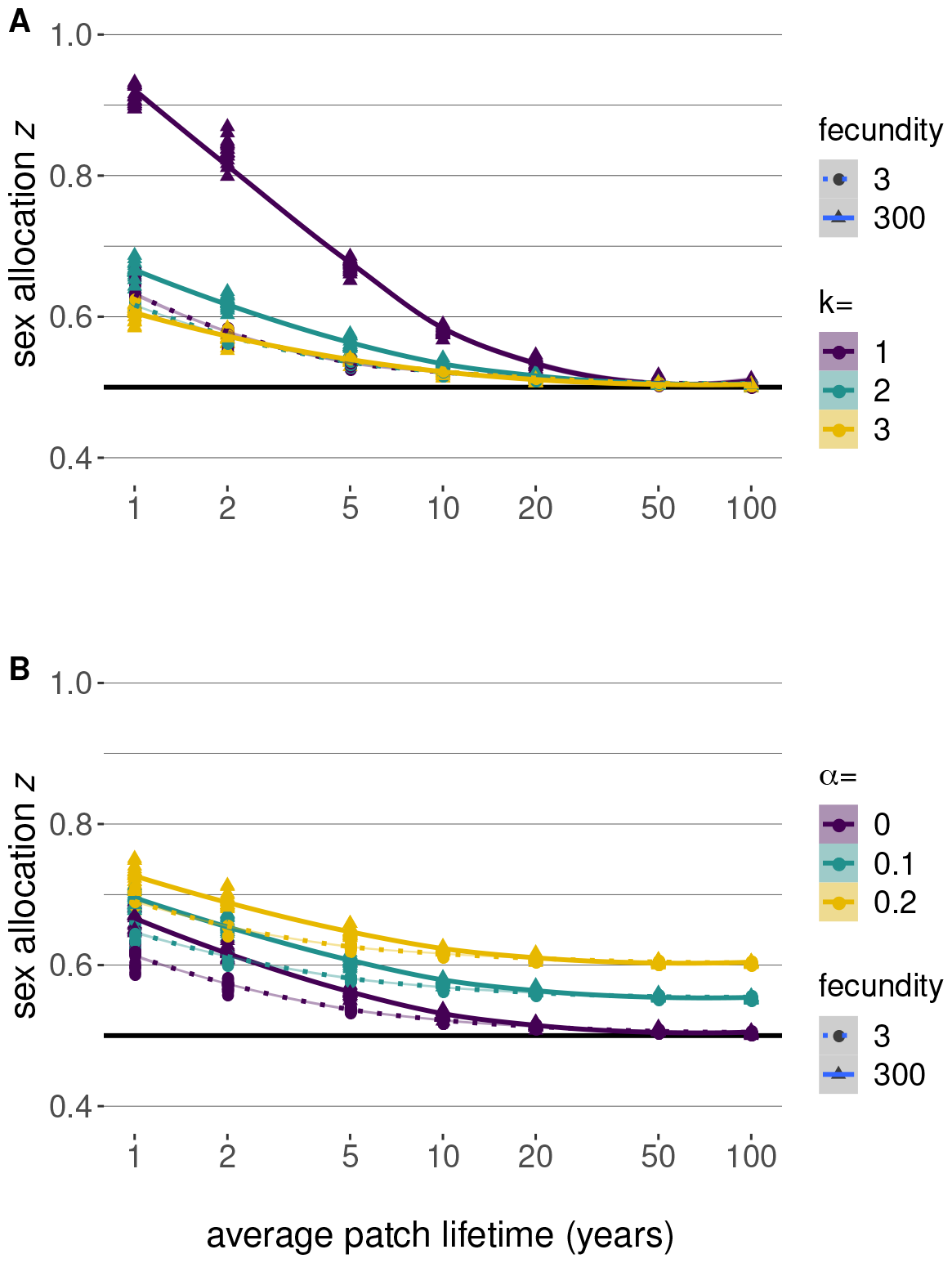
Effect of colonization bottlenecks and the rate of selfing on the female sex allocation in a metapopulation with migrant-pool colonization. A metapopulation of 1,000 demes of 100 individuals was simulated over 5,000 generations. 20 replicates were simulated for each parameter combination. In contrast with the model for the analytical calculations (Figure 1), the extinct demes are not assumed to become immediately filled to carrying capacity by a single round of reproduction among *k* colonizers. Fecundity is incorporated as the maximum number of seeds a hermaphrodite can produce. Two values for fecundity where explored here: fecundity = 3 (circles for individual simulations, dotted lines for the loess regression); and fecundity = 300 (triangles and solid lines). The extinction rate is transformed into the average lifetime of a deme. **A**. Seed dispersal was *d*_s_ = 0.05, pollen dispersal is *d*_p_ = 0 and *α* = 0. After extinction, demes were recolonized by *k* = 1 (purple), *k* = 2 (blue) or *k* = 3 (yellow) individuals. **B**. dispersal similar to panel A, but *α* = 0 (purple), *α* = 0.1 (blue) and *α* = 0.2 (yellow). *k* = 2 individuals.

**Figure S6:**
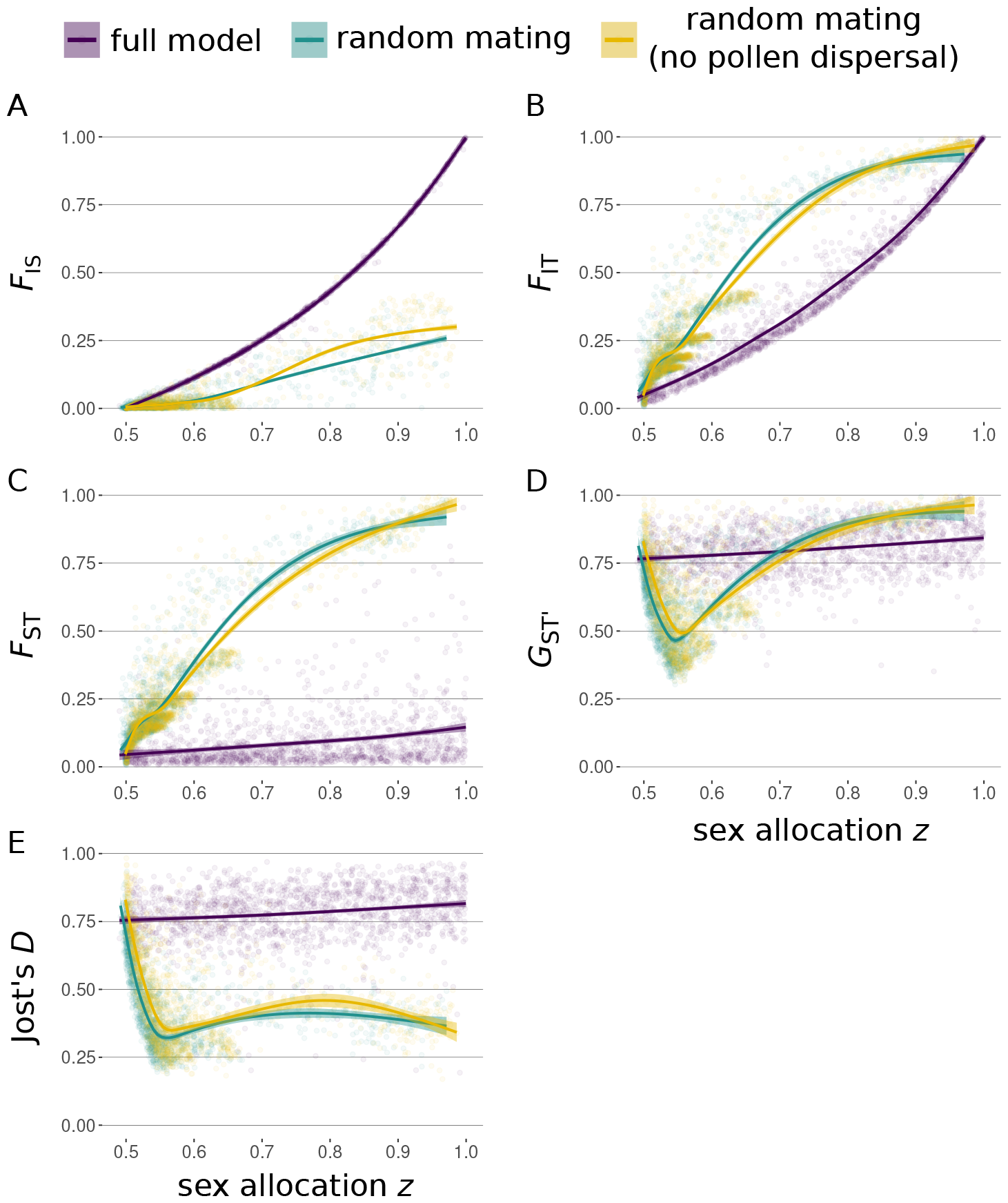
Association between sex allocation and *F*-statistics under a migrant-pool model A. Global *F*_IS_, **B**. *F*_IT_, **C**. *F*_ST_, **D**. *G*_*ST*_ ^*′*^, and **E**. Jost’s *D* are reported over 20 simulated unlinked loci, after 5,000 generations of evolution of a metapopulation composed of 1,000 demes, each with 1,000 hermaphrodites. 4,500 simulations in total were performed, distributed in three main conditions: full-model (purple) allows for seed dispersal, pollen dispersal, and self-fertilization; random mating (blue) allows for seed and pollen dispersal, but self-fertilization takes place only randomly at the time of sampling both parents; random mating without pollen dispersal (yellow) is similar to the previous condition but with *m*_p_ = 0. For each simulation, the parameters were randomly drawn in the following intervals: average deme lifetime in [1, 200] generations (i.e, extinction rate in [0.005, 0.5]); *m*_p_ and *m*_s_ in [0, 0.1]%; *α* in [0, 1]; *k* in [1, 6] individuals. In these simulations, the extinct and then recolonized demes are immediately filled by a round of reproduction among the *k* colonizers.

**Figure S7:**
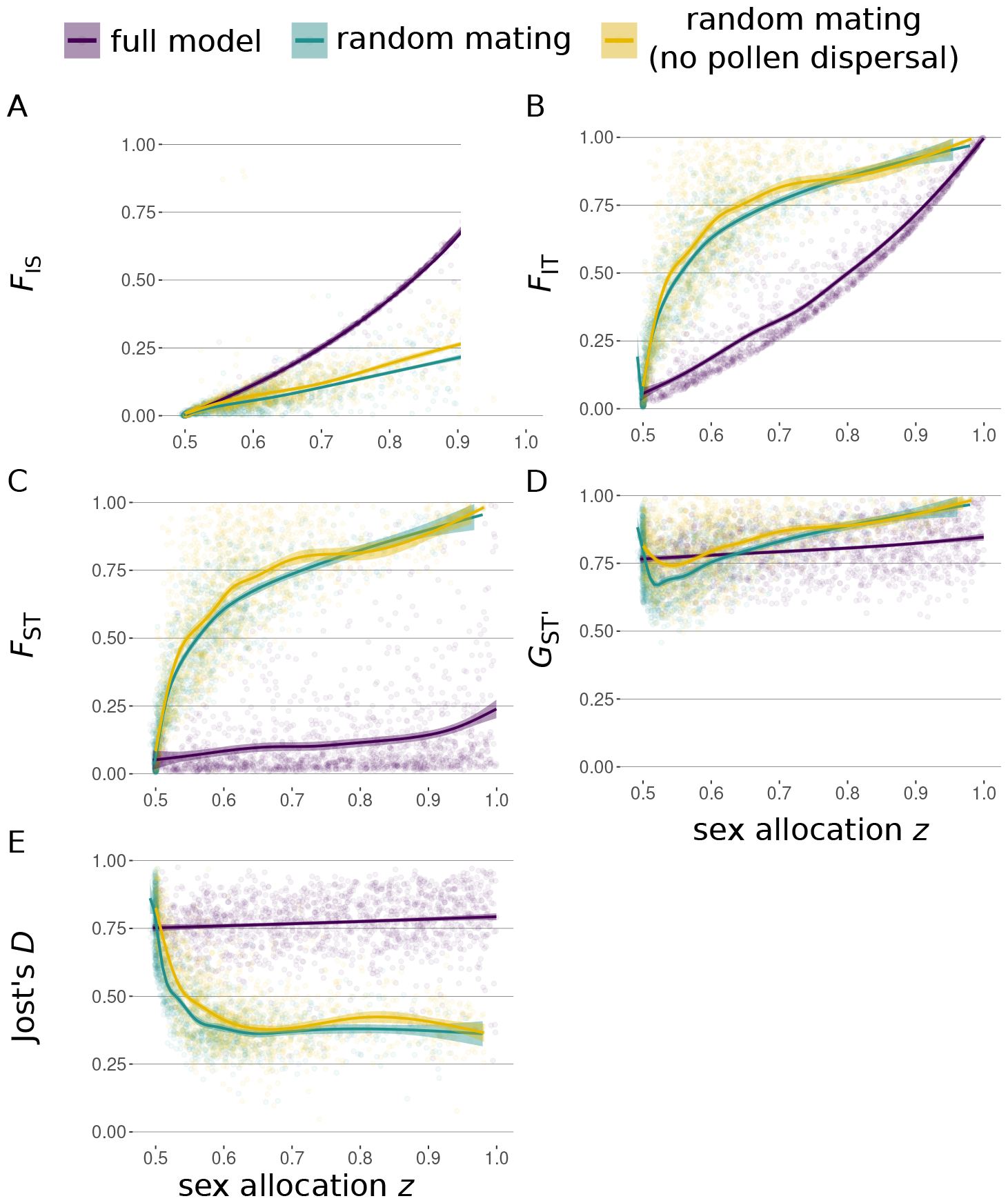
Association between sex allocation and *F*-statistics under a model of propagule-pool colonization. A. Global *F*_IS_, **B**. *F*_IT_, **C**. *F*_ST_, **D**. *G*_*ST*_ ^*′*^, and **E**. Jost’s *D* are reported over 20 simulated unlinked loci, after 5,000 generations of evolution of a metapopulation composed of 1,000 demes, each with 1,000 hermaphrodites. 4,500 simulations in total were performed, distributed in three main conditions: full-model (purple) allows for seed dispersal, pollen dispersal, and self-fertilization; random mating (blue) allows for seed and pollen dispersal, but self-fertilization takes place only randomly at the time of sampling both parents; random mating without pollen dispersal (yellow) is similar to the previous condition but with *m*_p_ = 0. For each simulation, the parameters were randomly drawn in the following intervals: average deme lifetime in [1, 200] generations (i.e, extinction rate in [0.005, 0.5]); *m*_p_ and *m*_s_ in [0, 0.1]%; *α* in [0, 1]; *k* in [1, 6] individuals. In these simulations, the extinct and then recolonized demes are immediately filled by a round of reproduction among the *k* colonizers.

## Notes

### Competing Interest Statement

The authors have declared no competing interest.

### Summary of Updates

The previous version of this article has been modified substantially through the inclusion of further analytical results and corresponding changes to the text. The article's conclusions remain largely unaltered.

